# Deep learning redesign of PETase for practical PET degrading applications

**DOI:** 10.1101/2021.10.10.463845

**Authors:** Hongyuan Lu, Daniel J. Diaz, Natalie J. Czarnecki, Congzhi Zhu, Wantae Kim, Raghav Shroff, Daniel J. Acosta, Brad Alexander, Hannah Cole, Yan Jessie Zhang, Nathaniel Lynd, Andrew D. Ellington, Hal S. Alper

**Affiliations:** McKetta Department of Chemical Engineering, The University of Texas at Austin, Austin, Texas 78712, United States; Department of Chemistry, The University of Texas at Austin, Austin, Texas 78712, United States; Department of Molecular Biosciences, The University of Texas at Austin, Austin, Texas 78712, United States

## Abstract

Plastic waste poses an ecological challenge^1^. While current plastic waste management largely relies on unsustainable, energy-intensive, or even hazardous physicochemical and mechanical processes, enzymatic degradation offers a green and sustainable route for plastic waste recycling^2^. Poly(ethylene terephthalate) (PET) has been extensively used in packaging and for the manufacture of fabrics and single-used containers, accounting for 12% of global solid waste^3^. The practical application of PET hydrolases has been hampered by their lack of robustness and the requirement for high processing temperatures. Here, we use a structure-based, deep learning algorithm to engineer an extremely robust and highly active PET hydrolase. Our best resulting mutant (FAST-PETase: <u>F</u>unctional, <u>A</u>ctive, <u>S</u>table, and <u>T</u>olerant PETase) exhibits superior PET-hydrolytic activity relative to both wild-type and engineered alternatives, (including a leaf-branch compost cutinase and its mutant^4^) and possesses enhanced thermostability and pH tolerance. We demonstrate that whole, untreated, post-consumer PET from 51 different plastic products can all be completely degraded by FAST-PETase within one week, and in as little as 24 hours at 50 °C. Finally, we demonstrate two paths for closed-loop PET recycling and valorization. First, we re-synthesize virgin PET from the monomers recovered after enzymatic depolymerization. Second, we enable *in situ* microbially-enabled valorization using a *Pseudomonas* strain together with FAST-PETase to degrade PET and utilize the evolved monomers as a carbon source for growth and polyhydroxyalkanoate production. Collectively, our results demonstrate the substantial improvements enabled by deep learning and a viable route for enzymatic plastic recycling at the industrial scale.

## Manuscript Text

Poly(ethylene terephthalate) (PET) composes 70% of synthetic textile fibers and 10% of non-fiber plastic packaging^1^, and correspondingly represents an enormous waste stream of single-use, manufactured materials. Yet, a circular carbon economy for PET is theoretically attainable through rapid enzymatic depolymerization followed by either chemical repolymerization or microbial upcycling/valorization into other products^5, 6^. However, all existing PET-hydrolyzing enzymes (PHEs) are limited in their capacity to either function within moderate pH/temperature ranges or directly utilize untreated post-consumer plastics. Such traits are essential for *in situ* depolymerization and for simplified, low-cost industrial-scale processes^7^. To overcome these limitations, we employed deep learning and protein engineering approaches to generate a PHE that has exceptionally high activity across a broad range of raw PET substrates (both model and actual post-consumer PET (pc-PET)), temperatures, and pH levels in a manner that out-performs all other known PHEs and rationally-derived mutants.

Enzymatic depolymerization of PET was first reported in 2005 and has been nascently demonstrated using 19 distinct PHEs derived from esterases, lipases, and cutinases^2, 7, 8^. However, the majority of these enzymes only show appreciable hydrolytic activity at high reaction temperatures (i.e. at or exceeding the PET glass transition temperature of ca. 70 °C) and with highly processed substrates. For example, an engineered leaf-branch compost cutinase (LCC) can degrade 90% of pretreated pc-PET within 10 hours at 72 °C and a pH of 8.0^4^. Most other PHEs similarly show poor activity at moderate temperatures^9^ and more neutral pH conditions^10^, greatly restricting *in situ* / microbially-enabled degradation solutions for PET waste. This limitation is of critical concern as 40% of uncollectable plastics reside in natural environments^11^. In addition, converting untreated post-consumer plastic waste at near ambient temperature would be preferable for industrial applications, whereas elevated temperatures and pre-treatment increase net operating costs.

While the PHE from the PET-assimilating bacterium *Ideonella sakaiensis*^9^ (PETase) can operate at ambient conditions, it is highly labile and loses activity even at 37 °C after 24 hours^12^, thereby limiting practical applications. Nonetheless, this mesophilic enzyme has previously seen attempts to enhance thermostability, robustness and function^12–18^. The most notable engineered PETase variants—ThermoPETase^12^ and DuraPETase^17^—were created through rational protein engineering and computational redesign strategies, respectively. Although the thermostability and catalytic activity of these two mutants were improved^12, 17^ under certain conditions, they nonetheless had overall lower PET-hydrolytic activity at mild temperatures.

We posited that highly focused protein engineering approaches such as those described above cannot take into account the evolutionary trade-off between overall stability and activity, and that a neutral, structure-based, deep learning neural network might generally improve enzyme function across all conditions. To this end, we employed our 3D self-supervised, convolutional neural network, MutCompute^19^ (Supplementary Information Fig. 1) to identify stabilizing mutations. This algorithm learns the local chemical microenvironments of amino acids based on training over 19,000 sequence-diverse protein structures from the Protein Data Bank and can readily predict positions within a protein where wild-type amino acids are not optimized for their local environments. We employed MutCompute to obtain a discrete probability distribution for the structural fit of all 20 canonical amino acids at every position in both wild-type PETase and ThermoPETase (crystal structures PDB: 5XJH and 6IJ6) (Supplementary Information Fig. 2), essentially carrying out a comprehensive scanning mutagenesis of the protein *in silico*. The predicted distributions were rendered onto the protein crystal structure (Fig. 1a) to identify positions where wild-type amino acid residues were ‘less fit’ than potential substitutions. Predictions were then ranked by predicted probabilities (fold-change of fit) (Fig. 1b; Supplementary Information Fig. 3). Using a stepwise combination strategy, a total of 159 single or multiple predicted mutations were generated in various PETase scaffolds. Variants exhibiting improved catalytic activity (as measured by esterase activity and plastic degradation rates) and thermostability (as measured by protein melting temperature (T_m_)) were characterized further. Amongst this set, four predicted mutations (S121E, N233K, R224Q and T140D) (Fig 1c) resulted in the highest improvements, both singly and in combination, and were selected for further assembly and analysis (see Additional Supplementary Discussion in Supplementary Information Fig. 4 for a further discussion of the mutant down-select). Encouragingly, two substitutions (S121E and T140D) were reported in the literature after our initial predictions, whereas the remaining residues are entirely unique, thus emphasizing the importance of a neutral, deep learning-based approach to identifying critical substitutions.

**Fig. 1.**
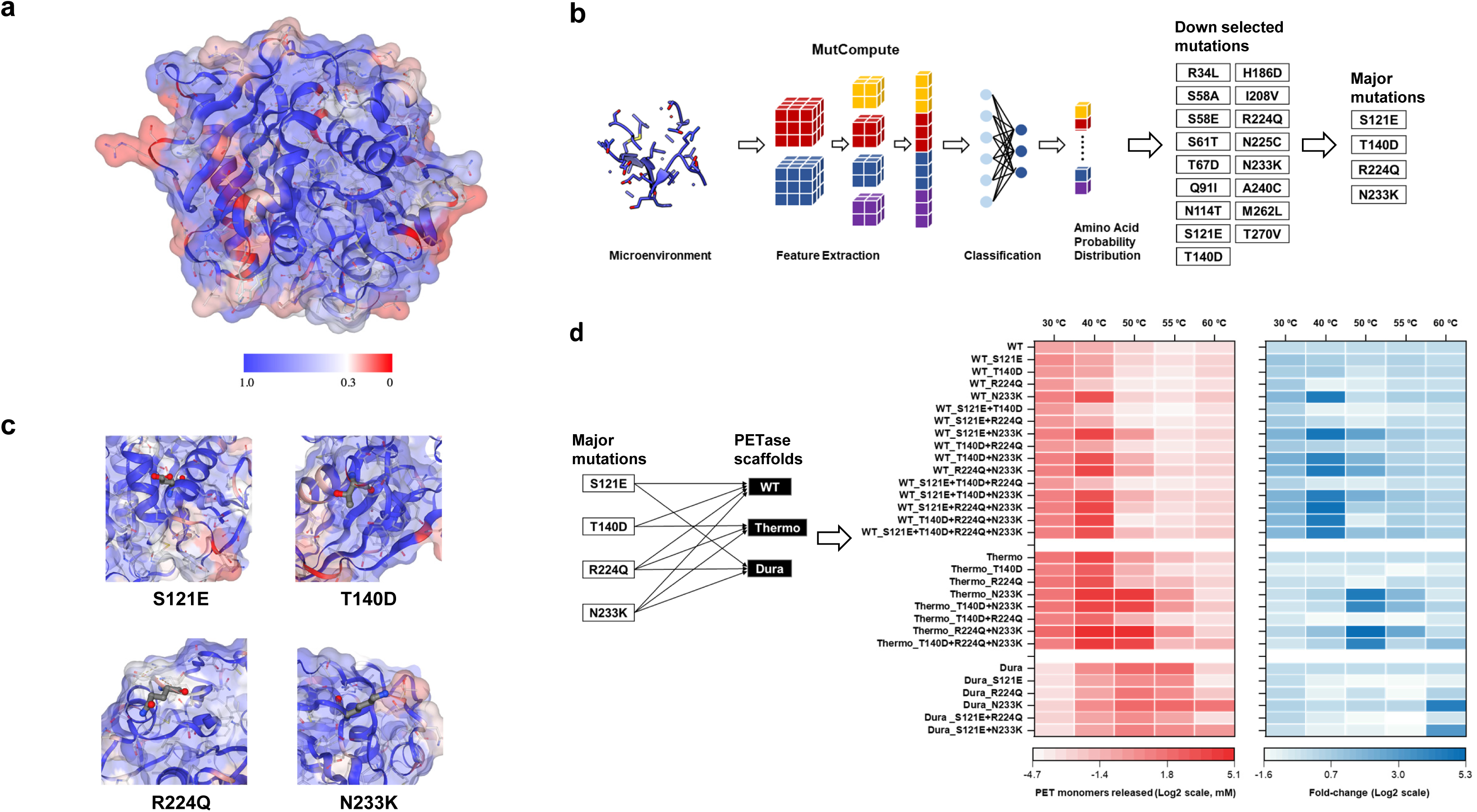
Machine-learning guided predictions improve enzyme performance across PETase scaffolds. **a.** Wild-type PETase protein structure rendered by the output of Mutcompute. Each amino acid residue was assessed by Mutcompute, resulting in a probability distribution reflecting the chemical congruency of each of the twenty amino acids with the neighboring chemical microenvironments. Residues assigned the low wild-type probability (disfavored) are red and the high wild-type probability (favored) are blue. Interactive visualizations of MutCompute are available at https://www.mutcompute.com/petase/5xjh and https://www.mutcompute.com/petase/6ij6. **b.** Using MutCompute with both wild-type PETase and ThermoPETase, two libraries of predictions were generated. To down-select mutations, the predictions were ranked by the fold change in the probabilities between the predicted and the wild-type amino acid. Using a stepwise combination strategy, 159 variants were generated by incorporating single or multiple predicted mutations into various PETase scaffolds. After experimental characterization, four predicted mutations (S121E, N233K, R224Q and T140D) resulted in the highest improvements both singly and in combination. **c.** Microenvironment of the four major mutations predicted by Mutcompute. **d.** The four major mutations were completely and combinatorially assembled across three PETase scaffolds: wild-type PETase (WT), ThermoPETase (Thermo), and DuraPETase (Dura). The red heatmap (left) shows the PET-hydrolytic activity of the resulting variants and the blue heatmap (right) shows the fold-change of activity over their respective scaffolds. PET-hydrolytic activity was evaluated by measuring the amount of PET monomers (the sum of TPA and MHET) released from hydrolyzing circular gf-PET film (6 mm in diameter, ∼11.4 mg) by the PETase variants after 96 hrs of incubation at temperature ranging from 30 to 60 °C. All measurements were conducted in triplicate (n=3).

We assembled all 29 possible combinations using these four mutations across three PETase scaffolds (wild-type PETase, ThermoPETase, and DuraPETase). Of note, two could not be purified using the DuraPETase background after multiple attempts. Thermostability analysis of the remaining 27 mutants indicated that 24 (ca. 89%) resulted in elevated T_m_ relative to their respective scaffolds (Supplementary Information Fig. 5). The highest change in thermostability from their respective PETase scaffolds were observed for variants PETase^N233K/T140D^ with a T_m_ of 58.1 °C (ΔT_m_=10 °C from WT PETase), ThermoPETase^N233K/R224Q^ with a T_m_ of 67.4 °C (ΔT_m_=9 °C from ThermoPETase), and DuraPETase^N233K^ with a T_m_ of 83.5 °C (ΔT_m_=5 °C from DuraPETase). The latter mutant represents the most thermostable PETase mutant reported to date. It was noted that the protein yield of all 27 variants was improved (up to 3.8-fold increase) compared with the parental scaffold, further underscoring the ability of Mutcompute to identify mutants of higher stability (Supplementary Information Fig. 6). The portability and combinatorial synergy of these mutations across scaffolds demonstrates the power of this neural network-based approach.

Next, we sought to evaluate the PET hydrolytic activity of these more stable variants across a range of temperatures from 30 to 60 °C using an amorphous PET film (gf-PET, from the supplier Goodfellow, PA, USA) commonly used in the literature^4^. This comparison immediately revealed that the machine-learning guided predictions greatly enhanced PET-hydrolytic activity and extended the range of working temperature in all scaffolds (Fig. 1d). In particular, PETase ^N233K/R224Q/S121E^ exhibited a 3.4-fold and 29-fold increase in PET-hydrolytic activity at 30 and 40 °C respectively, over wild-type PETase (Fig. 1d). Enzyme mutants based on the ThermoPETase scaffold showed an extended range of working temperature (30-60 ⁰C) and exhibited significantly higher activity than their counterparts. Within this set, the best variant from the ThermoPETase scaffold (containing N233K and R224Q on top of S121E), named FAST-PETase (<u>F</u>unctional, <u>A</u>ctive, <u>S</u>table, and <u>T</u>olerant PETase), showed 2.4-fold and 38-fold higher activity at 40 and 50 ⁰C, respectively compared to ThermoPETase alone (Fig. 1d). At 50 ⁰C, FAST-PETase displayed the highest overall degradation of all mutants and temperatures activity releasing 33.8 mM of PET monomers in 96 hours (Fig. 1d). The DuraPETase scaffold in general exhibited relatively low activity at mild temperatures (30–50 ⁰C), but improvements were nevertheless realized at higher temperatures (55–60 ⁰C) as demonstrated by the most thermostable PETase mutant-DuraPETase^N233K^ (Fig. 1d).

Crystal structure analysis of FAST-PETase at 1.44 Å resolution explains the enhanced stability through newly formed, favorable residue interactions (Fig. 2). The N233K mutation places a positively-charged lysine next to E204 and establishes an intramolecular salt bridge (Fig. 2f). The side chain of R224, when mutated to Gln, forms a hydrogen bond to the carbonyl group of S192 (Fig. 2d). Finally, the S121E mutation enables a new water-mediated hydrogen-bonding network with H186 and N172 (Fig. 2d).

**Fig. 2.**
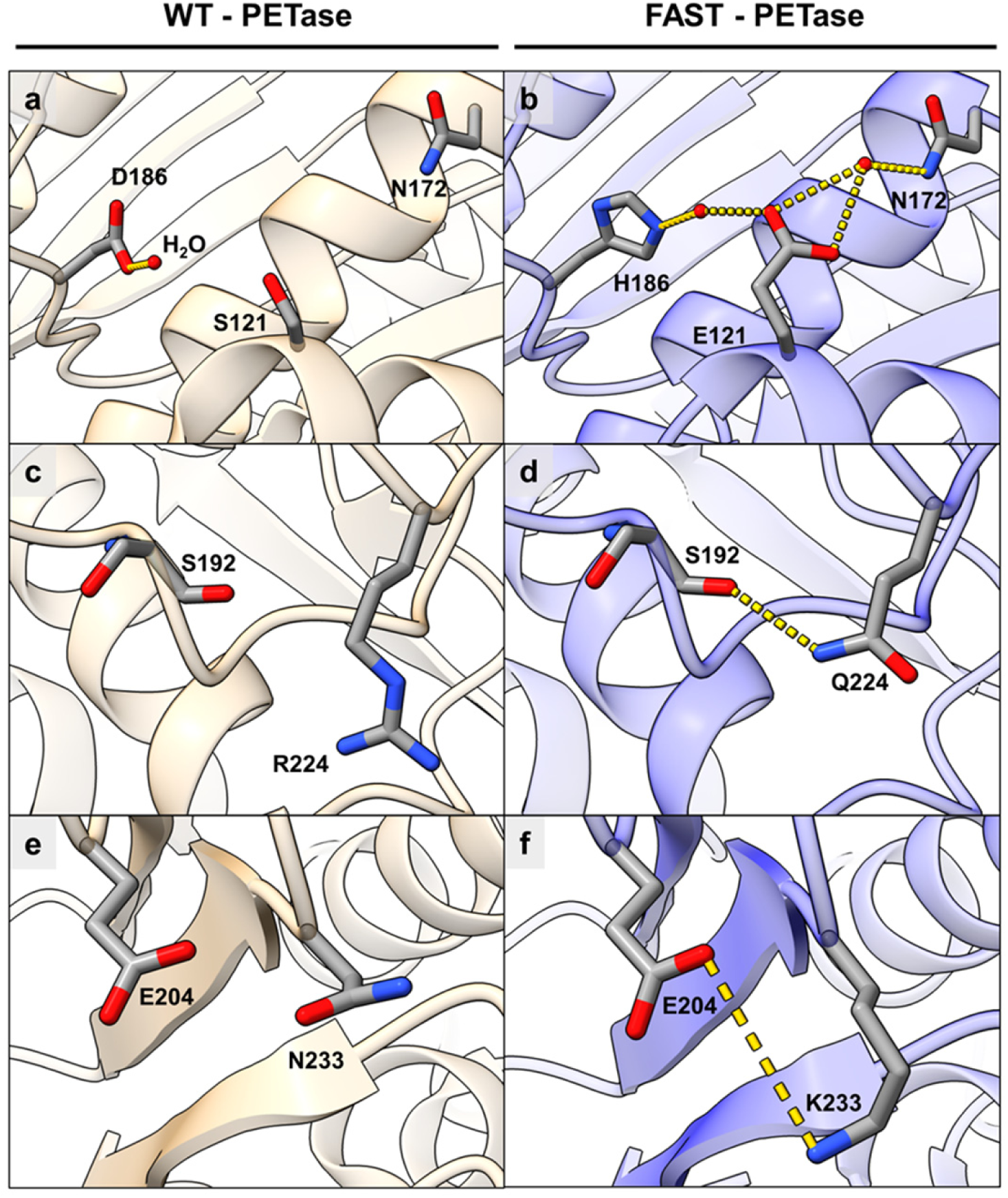
Predicted mutations from neural network algorithm stabilizes FAST-PETase. **a-f.** Structural comparison between (**a**, **c**, **e**) wild-type PETase (tan-colored stick model, PDB code: 5XJH) and (**b**, **d**, **f**) FAST-PETase (blue-colored stick model, PDB code: 7SH6) near the predicted mutation sites (S121E, R224Q, N233K respectively). Hydrogen bonding and salt bridge interactions are shown and highlighted as yellow dotted lines.

To evaluate the catalytic resilience of these mutants to environmental conditions, FAST-PETase were compared to previously reported wild-type and mutant PHEs including wild-type PETase, ThermoPETase, DuraPETase, LCC, the most active mutant LCC ^F243I/D238C/S283C/N246M^(ICCM) using gf-PET across a range of pH (6.5 – 8.0) and temperatures (30-40 ⁰C) (Supplementary Information Fig. 7). This comparative analysis demonstrated the unique catalytic capability of FAST-PETase to function at low pH levels and ambient temperature. Specifically, FAST-PETase outperformed other PHEs (including prior rational designs) at all pH conditions. Especially at pH 7, FAST-PETase exhibited activities that were 9.7 and 115 times as high as that of wild-type PETase at 30 and 40 ⁰C, respectively (Supplementary Information Fig. 7). This enzymatic performance makes FAST-PETase an excellent candidate for mild temperatures and moderate pH enzymatic degradation of PET seen in conditions of *in situ* plastic degradation.

Beyond model plastic substrates, it is critical to demonstrate the performance of PETase enzymes on raw, untreated pc-PET. Notably, unlike the gf-PET used above and throughout the literature, there is no singular pcPET substrate. To this end, we collected 51 samples of post-consumer plastic products used in the packaging of food, beverages, medications, office supplies, household goods and cosmetics available at local grocery store chains and treated this raw material enzymatically with FAST-PETase at 50 °C (Supplementary Information Fig. 8). Despite their heterogeneity including physical properties such as crystallinity, molecular weight, and thickness as well as different compositions including additives and plasticizers, hole-punched samples from this wide array of PET products were all fully degraded by FAST-PETase within one week and in as little as 24 hours (Fig. 3a). While thickness of the plastic did correlate with degradation time (as thickness and mass are related), neither this metric nor crystallinity or any other measured trait of PET alone determined overall degradation rates (Supplementary Information Fig. 9).

**Fig. 3.**
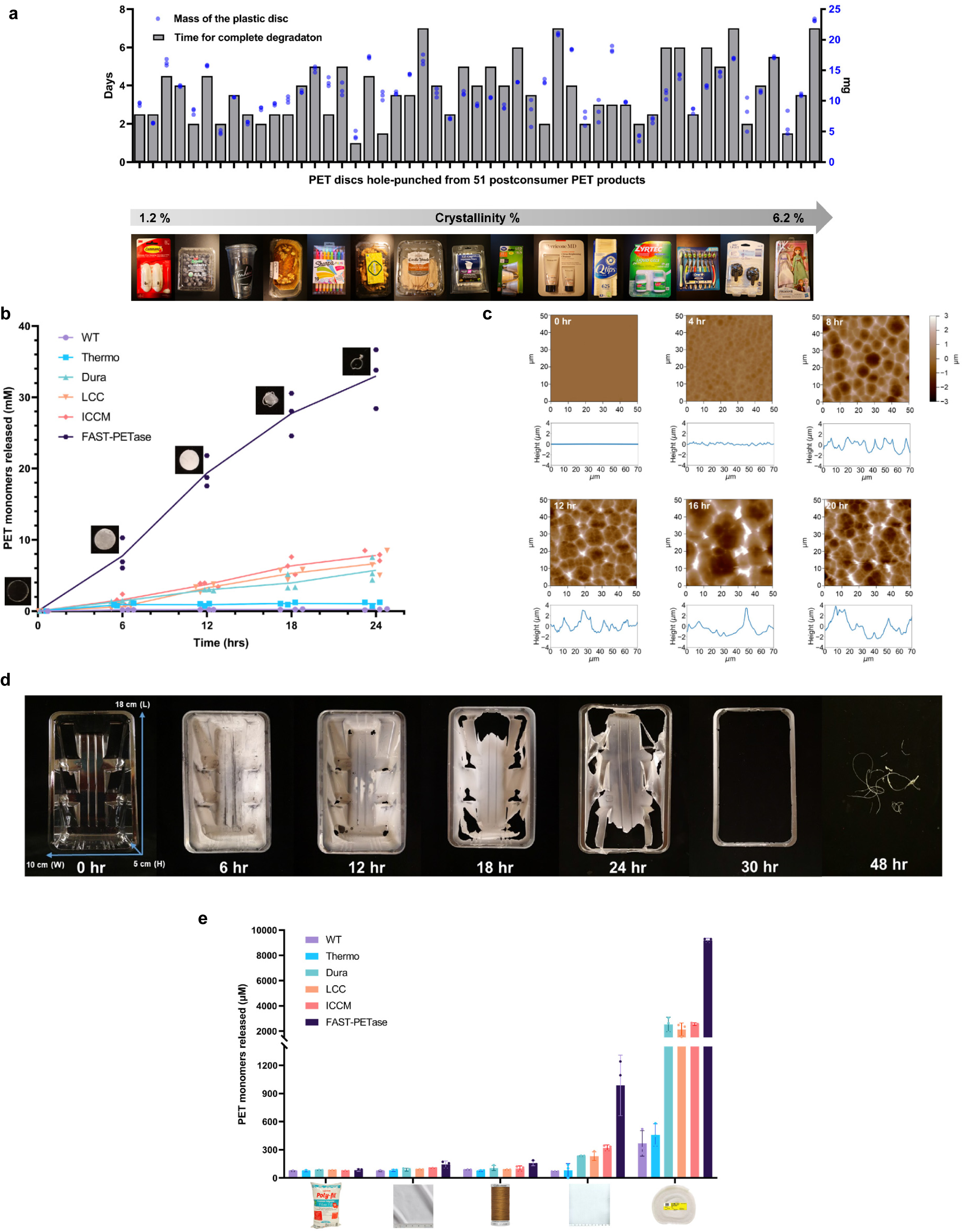
The superior performance of FAST-PETase in enzymatic depolymerization of post-consumer PET plastic and polyester products. **a.** Complete degradation of pc-PET films hole-punched from 51 post-consumer plastic products. **b.** Time-course of PET-hydrolytic activity of FAST-PETase, wild-type PETase (WT), ThermoPETase (Thermo), DuraPETase (Dura), LCC and ICCM at reaction temperature of 50 ⁰C. PET-hydrolytic activity was evaluated by measuring the amount of PET monomers (the sum of TPA and MHET) released from hydrolyzing pc-PET (Bean cake plastic container) film by the tested PHEs at various time points. KH_2_PO_4_-NaOH (pH 8) buffer was used for all enzymes shown in this figure. All measurements were conducted in triplicate (n=3). **c.** Atomic Force Microscopy images of pc-PET films following various exposure times with FAST-PETase. **d.** Complete degradation of large, untreated PET container with FAST-PETase at 50 ⁰C. **e.** Degradation of commercial polyester products with FAST-PETase, wild-type PETase (WT), ThermoPETase (Thermo), DuraPETase (Dura), LCC and ICCM at 50 ⁰C. All measurements were conducted in triplicate (n=3).

Among the post-consumer products tested above, we further evaluated the sample from a bean cake container that was completely degraded by FAST-PETase within 24 hrs at 50 ⁰C. A time-course analysis (Fig. 3b) revealed that the degradation of this pc-PET film exhibited an almost linear decay rate using FAST-PETase in terms of the total PET monomers released. Concomitantly, degradation of the pc-PET film by FAST-PETase brought an increase in the crystallinity from 1.2 % to 7.7% over 24 hrs (Supplementary Information Fig. 10). Atomic Force Microscopy (Fig. 3c) as well as Scanning Electron Microscopy (Supplementary Information Fig. 11) further showed that the reaction progression of FAST-PETase as it produced increasingly deeper and larger holes in the pc-PET surface resulting in increased surface roughness (and visible opaqueness) over reaction time (Supplementary Information Fig. 12). In contrast, the PET-hydrolytic activity of wild-type PETase, ThermoPETase, DuraPETase, LCC and ICCM toward this pc-PET was substantially lower (3.2 to 141.6-fold) than that of FAST-PETase under the same conditions (Fig. 3b). Interestingly, even at their previously reported optimal reaction temperature of 72 ⁰C ^4^, the activity of LCC and ICCM was still 4.9-fold and 1.5-fold lower than that of FAST-PETase at 50 ⁰C. Further experimental analysis (Supplementary Information Fig. 13) indicated that LCC and ICCM exhibited their highest degradation rate against this pc-PET film at 60 ⁰C. However, even at 60⁰C, the activity of LCC and ICCM was still lower than that of FAST-PETase at 50 ⁰C. Moreover, we demonstrate that the depolymerization process with FAST-PETase is easily scalable to large, untreated pieces of plastic (in this case, 6.4 g rather than 11 mg) simply by increasing net reaction volumes (Fig. 3d). Given these results, FAST-PETase can serve as a promising biocatalyst for the enzyme-based platform aimed at recycling raw, untreated PET waste, with advantages of lower operating cost and higher degradation efficiency of pc-PET, in contrast to ICCM that requires a higher reaction temperature.

Beyond packaging materials, PET is used heavily in the synthetic textile industry. To this end, we evaluated the potential application of FAST-PETase to partially degrade commercial polyester products. Five different commercial polyester products were treated with FAST-PETase at 50 ⁰C, releasing higher amounts of terephthalic acid (TPA) and Mono-(2-hydroxyethyl)terephthalate (MHET) relative to that of the samples treated with other PHEs (Fig. 3e). This indicates that FAST-PETase can potentially be used for rapid and efficient degradation of the PET fragments embedded in textile fabrics, providing a potential route for recovering PET monomers from commercial polyester products and reducing the leaching of microfibers into the environment.

Given the high activity of this FAST-PETase mutant at ambient temperatures and pH conditions, we hypothesized that this enzyme would be suitable for various enzymatic-microbial and enzymatic-chemical processing of PET. In this regard, PET depolymerization is only half of the circular plastic economy and we demonstrate here the compatibility of FAST-PETase with both chemical and biological recycling/upcycling applications to close the cycle. First, we demonstrate a closed-cycle PET re-constitution by first depolymerizing a tinted post-consumer plastic waste utilizing FAST-PETase and subsequently recovering monomers. TPA was recovered from the degradation solution with a yield of 96.8% and with a purity of over 99%. We then regenerate virgin PET directly from the degradation solution using chemical polymerization (Fig. 4a). A complete cycle of degradation to re-polymerization can be accomplished in as little as a few days (Supplementary Information Fig. 14). These results demonstrate the feasibility of a closed-loop enzymatic/chemical recycling process to generate a clear, virgin PET film from non-petroleum resources. Moreover, this workflow bypasses the challenges of recycling mixed-color PET products.

**Fig. 4.**
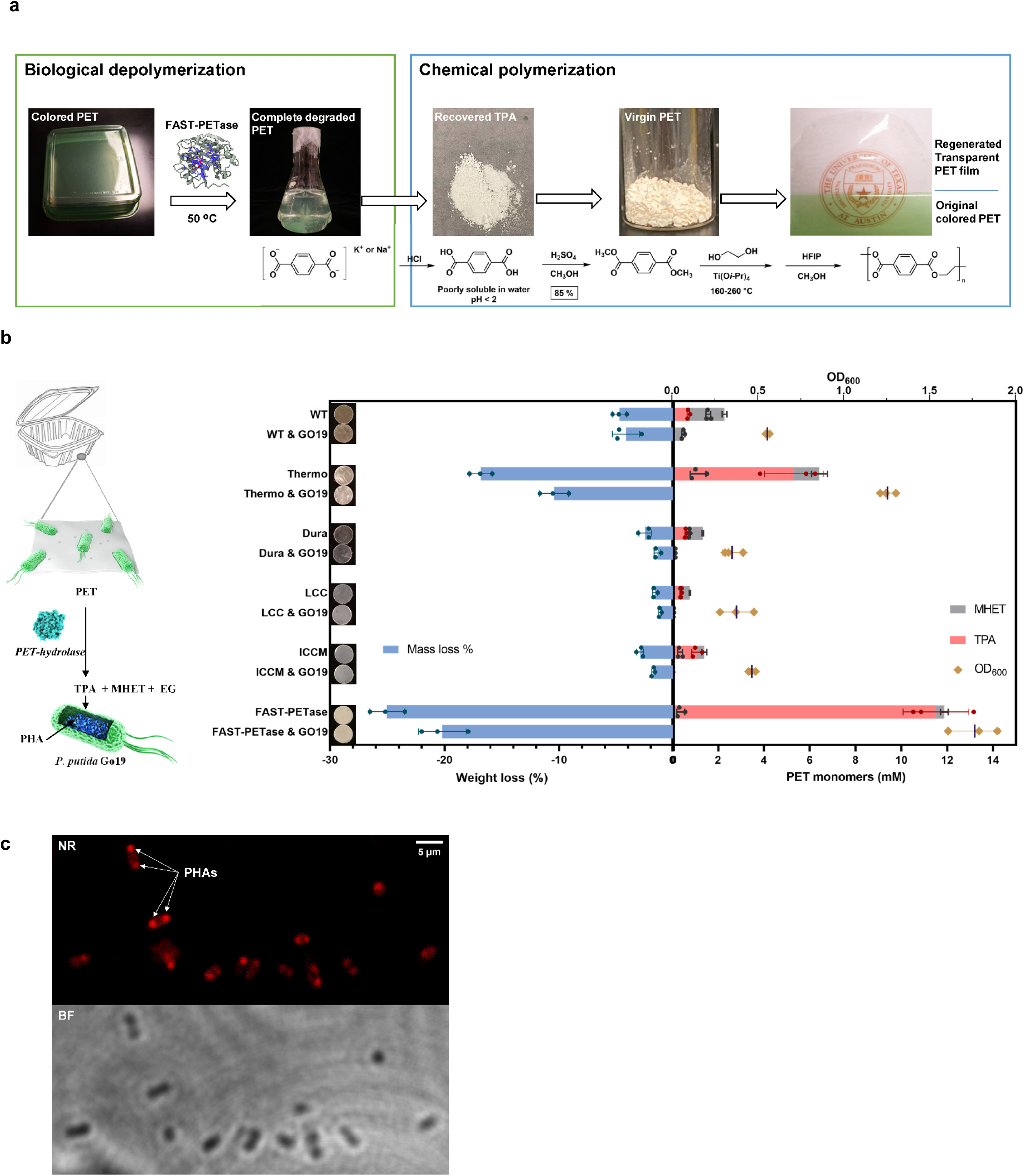
Applications of FAST-PETase in enzymatic-chemical recycling of PET and *in situ* depolymerization. **a.** Schematic of the closed-loop PET recycling process incorporating post-consumer colored plastic waste depolymerization by FAST-PETase and chemical polymerization. The crystallinity of the regenerated PET was determined as 58 % by Differential Scanning Calorimetry. The molecular weights (Mn, Mw), polydispersity indices (Ð) of the regenerated PET were determined as Mn = 16.4 kg/mol, Mw = 45.9 kg/mol, Ð = 2.80 by Gel Permeation Chromatography. **b.** Simultaneous process combining *P. putida* Go19 and exogenous PHEs: FAST-PETase, wild-type PETase (WT), ThermoPETase (Thermo), DuraPETase (Dura), LCC and ICCM. PHE (FAST-PETase/WT/Thermo/Dura/LCC/ICCM) & Go19 represents the simultaneous process of a PHE with *P. putida* Go 19, whereas PHE alone represents the control condition where the enzyme is presented without *P. putida* Go 19. PET monomers, mass loss of the pc-PET films, and cell density of *P. putida* Go19 were measured after 96 hrs of incubation. All measurements were conducted in triplicate (n=3). EG represents ethylene glycol. **c.** Bright field (BF) microscopy of *P. putida* Go19 cells after growth with pc-PET film for 96 hrs, and fluorescent microscopic observation of the intracellular PHAs granulate via Nile Red (NR) staining of same cells.

Second, we sought to utilize the degradation capability of FAST-PETase at ambient temperature to enable direct depolymerization and microbial valorization of monomers. To this end, we evaluated a simultaneous biodegradation scheme using FAST-PETase to demonstrate that this mutant enzyme is microbe-compatible. In particular, a soil bacteria *Pseudomonas putida* Go19^20, 21^ capable of naturally utilizing TPA as a carbon and energy source and capable of producing polyhydroxyalkanoates (PHAs) was employed. Initially, we sought to combine exogenous FAST-PETase with this host to explore the possibility of simultaneous PET depolymerization and fermentation. *P. putida* Go19 was inoculated into a minimal medium supplemented with an unpretreated pc-PET film absent of any other carbon source. Upon adding 200 nM of purified FAST-PETase to the culture medium, growth of *P. putida* Go19 was observed concomitant with the degraded pc-PET film which displayed opacity and lost 20.2 ± 2.1% of its initial weight (Fig. 4b) after 4 days. Through this experiment, we observed that the TPA liberated from the hydrolysis of pc-PET film by FAST-PETase was consumed by the *P. putida* Go19 for growth (Fig. 4b) and PHAs accumulation (Fig. 4c). In contrast, when wild-type PETase, ThermoPETase, DuraPETase, LCC, or ICCM was used as the catalyst in such process, the cell density of *P. putida* Go 19 and the weight loss of the pc-PET film were all significantly lower than when FAST-PETase was used (Fig. 4b). These results demonstrated that FAST-PETase exhibited the highest PET-hydrolytic activity under cell-growth compatible conditions when compared with other PHEs tested. This demonstration represents the first simultaneous bioprocess that integrates enzymatic PET depolymerization and TPA conversion to PHAs at ambient temperatures and neutral pH.

In conclusion, this work utilized a structure-based deep learning model to identify portable substitutions that imparted improved stability and function across a variety of PETase scaffolds. The best variant, FAST-PETase, exhibits superior activity over a wide range of temperatures (30–50 ⁰C), and exceptional compatibility with cell-growth conditions. We demonstrate this capacity via the rapid, efficient, and complete degradation of bulk, untreated pc-PET waste and a reduction of PET fragments embedded in textile fabrics. The properties of this variant are ultimately suitable for both low-cost industrial recycling as well as for *in situ* plastic degradation applications, as demonstrated by simultaneous bioprocessing with *P. putida* Go 19. Collectively, these results demonstrate the potential for structure-based deep learning in protein engineering and the opportunities for converting mesophilic enzyme scaffolds into broad-range biocatalysts for a cyclic plastic economy.

## Acknowledgements

This work was funded under research agreement EM10480.26 / UTA16-000509 between the ExxonMobil Research and Engineering Company and The University of Texas at Austin. Sequencing was conducted at the Genomic Sequencing and Analysis Facility and Scanning Electron Microscopic analysis was conducted at the Microscopy and Imaging Facility at the UT Austin Center for Biomedical Research Support. We acknowledge Dr. Sapun Parekh (The University of Texas at Austin) for access to his microscopic imaging facility, as well as Yujen Wang for his assistance in taking microscopic fluorescent images.

## Accession number

Coordinates for the FAST-PETase structure have been deposited into the Protein Data Bank with accession code 7SH6.

## Supplementary Methods

**Supplementary Methods 1:** Description of methods

### Convolutional Neural Network (CNN) Model

MutCompute^19^ is a 3D CNN model where the architecture consists of nine layers divided into two blocks: 1) feature extraction and 2) classification. The feature extraction block consisted of six layers: two pairs of 3D convolutional layers followed by a dimension reduction max pooling layer after each pair. The first pair of convolutional layers used filters of size 3x3x3 and the second pair had filters of size 2x2x2. Additionally, the Rectified Linearity Unit function (Relu) was applied to the output of each of the four convolutional layers. The final feature maps generated by the feature extraction block had dimensions of size 400x3x3x3. These feature maps were flattened into a 1D vector of size 10,800 before being passed to the classification block. The classification block consisted of three fully connected dense layers given dropout rates of 0.5, 0.2, and 0, respectively. Like the feature extraction block, the output of the first two dense layers was transformed by the Relu function. To obtain a vector of 20 probability scores representing the network prediction for each of the amino acids, we applied a softmax activation function to the output of the third dense layer. MutCompute is available as a machine learning as a service (MLaaS) on https://mutcompute.com.

### MutCompute Predictions

MutCompute predictions were obtained by running the wild type PETase (pdb id: 5xjh) and thermopetase (pdb id: 6ij6) through our MLaaS at https://mutcompute.com. Residues were filtered by sorting by the probability assigned to the wild type amino acid. We filtered 34 and 39 residues from the wild type PETase and Thermopetase crystal structures, respectively. From these filtered residues, we prioritized our experimental mutagenesis by selecting the 10 residues from each crystal structure with the highest log2 fold change between the predicted amino acid and wild type amino acid probabilities. Prior to experimentation, selected residues were visualized with Mutcompute-View, which is built on top of NGLViewer (https://github.com/nglviewer/ngl), to ensure three things: 1) the prediction was chemically sound and not due to a crystal or model artifact, 2) we were avoiding the active site and binding pocket and 3) avoiding epistatic interactions between predictions and instead targeting predicted “instability hotspots”. MutCompute-View has been made publicly available at https://mutcompute.com/view.

### Protein crystallization, X-ray diffraction data collection, data processing, and model refinement

To identify crystallization conditions for FAST-PETase, a sample containing 6 mg/mL purified FAST-PETase was screened in sparse-matrix screening using Phoenix robotic system (Art Robinson). The initial hits were identified with the rod-shaped crystals formed after incubating screening plates at 25 °C for three days. The crystallization was optimized as 0.1 M bis-TRIS, 2.0 M Ammonium Sulfate, pH 5.5 setup as sitting-drop vapor diffusion.

Individual FAST-PETase crystals were flash-frozen directly in liquid nitrogen after cryoprotected with 30 % (v/v) glycerol. X-ray diffraction data were collected at 23-ID-B beamline in Advance Photon Source (Lemont, IL). The X-ray diffraction pattern was processed to 1.44Å resolution for FAST-PETase crystals using HKL2000^22^. The structure was solved by molecular replacement with ThermoPETase structure as the initial search model (PDB code 6IJ6^12^). The molecular replacement solution for FAST-PETase structure was iteratively built and refined using Coot^23^ and Phenix^24^ refinement package. Procheck and MolProbity evaluated the quality of the finalized FAST-PETase structure. The final statistics for data collection and structural determination are shown in **Supplementary Information Fig. 18**.

### Cloning

Genes encoding *Ideonella sakaiensis* 201-F6 *Is*PETase (wild-type PETase) (Genbank: BBYR01000074, *ISF6_4831*), its mutants—PETase S121E/D186H/R280A (ThermoPETase^12^) and PETaseS214H/I168R/W159H/S188Q/R280A/A180I/G165A/Q119Y/L117F/T140D (DuraPETase^17^), _leaf-branch compost_ cutinase (LCC) (Genbank: AEV21261) and its mutant LCC ^F243I/D238C/S283C/N246M^(ICCM)^4^ were commercially synthesized for cloning. To enable extracellular expression of PETase and its mutants in *Pseudomonas putida* KT2440 (ATCC 47054), the nucleotide sequence of the signal peptide— SPpstu (21 amino acids) from maltotetraose-forming amylase of *Pseudomonas stutzeri* MO-19^25^ was used to substitute the original signal peptide sequence (first 27 amino acids) of wild-type PETase. The wild-type PETase and its mutants’ genes with SPpstu presented at the N-terminus were amplified by polymerase chain reaction (PCR) using the synthetic genes as a template. Subsequently, using Gibson Assembly method, DNA fragments encoding wild-type PETase, ThermoPETase and DuraPETase were respectively subcloned into a modified **pBTK552** vector^26^ where the antibiotic resistance marker was swapped from spectinomycin to kanamycin resistance gene and a C-terminal hexa-histidine tag-coding sequence was added. To enable intracellular expression of LCC and ICCM in *Escherichia coli* (DE3) (New England Biolabs, Ipswich, MA), the nucleotide sequence encoding the original signal peptide was removed from the synthetic genes. The LCC and ICCM genes without signal peptide sequence were amplified by PCR using the synthetic genes as a template. Subsequently, using Gibson Assembly method, the DNA fragments encoding LCC and ICCM were respectively subcloned into a commercial vector-pET-21b (Novagen, San Diego, CA), which carry a C-terminal hexa-histidine tag-coding sequence. The electrocompetent cell *E. coli* DH10β was transformed with the Gibson Assembly product by following a standard electroporation protocol. The resultant expression plasmid DNA was extracted from the overnight culture of the cloning host by using the QIAprep Spin Miniprep kit (Qiagen, Valencia, CA). The DNA sequence of extracted plasmid was verified by Sanger sequencing. List of nucleotide sequences and expressed amino acid sequences of the genes used in the study is provided in S**upplementary method 2**.

### Variant construction

Variants of wild-type PETase, ThermoPETase and DuraPETase were generated through site-directed mutagenesis using the PCR method described in the Q5^®^ Site-Directed Mutagenesis Kit E0552S (New England Biolabs, Ipswich, MA). The constructed plasmids carrying wild-type PETase, ThermoPETase and DuraPETase genes were used as the templates for mutagenesis PCR reaction. The corresponding primer sequences and annealing temperature were designed and generated by using the NEBaseChanger™ tool. Stellar™ Competent Cells (Clontech Laboratories, Mountain View, CA) were used as the cloning host and transformed with the ligated plasmids using the heat-shock method provided in the manufacturer’s instruction. Plasmid extraction and sequencing for the variants were performed under the same conditions as described for plasmids carrying wild-type PETase, ThermoPETase and DuraPETase genes.

### Protein expression and purification

For extracellular protein expression of wild-type PETase, ThermoPETase, DuraPETase, and their variants, *P. putida* KT2440 was used as the expression host and its electrocompetent cell was transformed with the corresponding constructed plasmids. For intracellular protein expression of LCC and ICCM, *E. coli BL21* (DE3) was used as the expression host and its electrocompetent cell was transformed with the corresponding constructed plasmids. A single colony of an *P. putida* KT2440 or *E. coli BL21* (DE3) strain harboring one of the constructed plasmids was inoculated into 2 mL of Luria Bertani broth (LB) medium with 50 µg/mL kanamycin and grown overnight at 37 °C/225 rpm. The overnight-grown culture (using 150 µl) was scaled up with 1000-fold dilution in a 500-mL triple baffled shake flask and grown to a cell density of 0.8 (optical density [OD_600_])at 37 °C/225 rpm. Protein expression was induced by adding 0.2 mM of isopropyl β-D-1-thiogalactopyranoside (IPTG) and cells were cultured for 24 hrs at 30 °C/225 rpm. For isolation of the extracellularly expressed PETase enzymes, the induced cell culture was centrifuged at 14,000 g and 4 ⁰C for 20 mins to obtain the supernatant that accommodates secretory protein. For isolation of the intracellularly expressed LCC and ICCM, the induced cell culture was harvested by centrifugation at 4,000 g and 4 ⁰C for 20 mins. Cell pellets were then resuspended in 25 mL of Dulbecco’s Phosphate Buffered Saline (DPBS) (Thermo Fisher Scientific, Waltham, MA) pH 7.0 buffer containing 10 mM imidazole, 1 g/L of lysozyme and 5 µl of Pierce™ Universal Nuclease (Thermo Fisher Scientific, Waltham, MA), followed by mixing on a rocker for 20 mins at 4 ⁰C. Subsequently, cells were lysed by sonication and the resulting cell lysate was centrifuged at 14,000 g and 4 ⁰C for 20 mins to obtain the supernatant that contains soluble proteins. Target proteins from the above two type of supernatants were both purified by HisPur™ Ni-NTA Resin (Thermo Fisher Scientific, Waltham, MA) according to the manufacturer’s instruction. Desalting of the protein eluent was carried out by using Sephadex G-25 PD-10 columns (GE Healthcare, Piscataway, NJ) according to the manufacturer’s instruction. All purification and desalting steps were performed at 4° C in a cold room. Afterwards, the purified protein was concentrated to a final volume of 1 mL by the 50 mL, 10KDa cut-off Amicon® Ultra Centrifugal Filters device (EMD Millipore Corporation, Billerica, MA) and preserved in DPBS (pH 7.0). The protein concentration was then determined by using the Coomassie Plus Bradford Assay kit (Thermo Fisher Scientific) and the Infinite M200 PRO microplate reader (Tecan, Männedorf, Switzerland) to measure the absorbance of assay mixtures. The presence and purity of the purified proteins were assessed by sodium dodecyl sulfate-polyacrylamide gel electrophoresis.

### In Vitro Analysis of PET hydrolytic activity using commercial Goodfellow PET film (gf-PET)

To evaluate the PET hydrolytic activity of PETase and its variants, the homogenous amorphous gf-PET film (Goodfellow, U.S. 577-529-50; specification: 1.3-1.4 g cm^-3^ density, 1.58-1.64 refractive index, 100×10^-13^ cm^3^. cm cm^-2^ s^-1^ Pa^-1^ permeability to water @25°C, 20-80 x10^-6^ K^-1^ coefficient of thermal expansion, 0.13-0.15 W m-1 K-1 @23°C thermal conductivity) was used as the substrate for degradation assays with the purified PETase enzyme and its variants. The gf-PET film was prepared in a circular form (6 mm in diameter, ∼11.4 mg) and was washed three times with 1 % SDS, 20 % ethanol, and deionized water before usage. Subsequently, the gf-PET film was put into a glass test tube and fully submerged in 600 µl of 100 mM KH_2_PO_4_-NaOH buffer (pH 8.0) with 200 nM purified enzyme. The test tube was tightly capped and wrapped with parafilm to minimize volatilization. The reaction mixture was then incubated at 30/40/50/55/60 °C for 96 hrs. Followed by removing the enzyme-treated gf-PET film from the reaction mixture, the enzyme reaction was terminated by heating at 85 °C for 20 mins. The reaction mixture samples were then centrifuged at 10,000 × g for 5 mins. The supernatant of each sample was further analyzed by High-performance liquid chromatography (HPLC) for quantifying PET monomers released from the PET depolymerization.

To compare the PET hydrolytic activity of FAST-PETase with wild-type PETase, ThermoPETase, DuraPETase, LCC, and ICCM across a range of pH (6.5 – 8.0) at 30 ⁰C and 40 ⁰C, similar experimental setup was used. The circular gf-PET film (6 mm diameter, ∼11.4 mg) was used as the substrate. The enzyme reactions were performed with 200 nM purified enzyme in 600 µl of 100 mM KH_2_PO_4_-NaOH buffer with various pH (6.5, 7.0, 7.5 or 8.0) at 30 ⁰C and 40 ⁰C. After incubating the enzyme reaction for 96 hrs, the supernatant of the reaction mixture was also analyzed by HPLC for quantifying PET monomers released from the PET depolymerization.

### Degradation of post-consumer PET (pc-PET) plastics

Hole-punched pc-PET films (6 mm in diameter) from 51 post-consumer plastic products were serially treated by 200 nM FAST-PETase in 600 µl of 100 mM KH_2_PO_4_-NaOH buffer (pH 8.0) at 50 ⁰C. Fresh enzyme solution was replenished every 24 hrs to maximize enzymatic degradation rate for degrading the 51 samples of various pc-PET films.

The time-course analysis of degrading a pc-PET film (Bean cake container) by various PET-hydrolyzing enzymes (PHEs) was conducted at 50 ⁰C. The reactions were performed with 200 nM enzyme in 600 µl of 100 mM KH_2_PO_4_-NaOH buffer (pH 8.0) and terminated at 6 hrs, 12 hrs, 18 hrs or 24 hrs for quantifying total PET monomers released at each time point.

A large, untreated, and transparent pc-PET (Bean cake container, 6.4 g) was treated by 200 nM of FAST-PETase in 100 mM of KH2PO_4_-NaOH buffer (pH 8.0) at 50 ⁰C. The whole piece of transparent PET container was fully submerged in 2.5 L of enzyme solution and completely degraded after 48 hrs.

### Degradation of polyester product

Five different commercial polyester products were cut into small pieces and used as the substrates that were fully submerged in 600 µl of 100 mM KH_2_PO_4_-NaOH buffer (pH 8.0) with 200 nM purified enzyme. Enzyme treatment on these five polyester products was conducted at 50 ⁰C. After 24 hrs of incubation, the reaction was terminated. The enzyme degradation solution was used to determine how much PET monomers were released from hydrolyzing these polyester products by various PHEs.

### Degradation of large, untreated pc-PET container and regeneration of virgin PET

A large, untreated, and green colored pc-PET container was cut into small rectangular flakes (*ca.* 1cm ×3 cm). 3.0 g of the colored pc-PET flakes was serially treated by 200 nM FAST-PETase in 100 mM KH_2_PO_4_-NaOH buffer (pH 8.0) at 50 ⁰C. 200 nM of fresh enzyme was added to the degradation solution every 24 hrs to maximize enzymatic degradation rate. All the colored pc-PET fragments were completely degraded after 6 days.

Upon completion of degradation, the enzyme degradation solution was filtered, and the filtrate was collected for regeneration. The pH of the filtrate was adjusted to pH 11 with NaOH to hydrolyze remaining MHET. The solution was stirred at room temperature for 4 hrs to complete the hydrolysis. The pH of the solution was subsequently adjusted to 2 with 37% HCl. The precipitate was filtered and washed several times with deionized water and dried under vacuum overnight. 4.3 g of TPA was collected and used in the next step without further purification. ^1^H NMR (400 MHz, *d*_6_-DMSO): 8.03 ppm (s, 4H).

To a suspension of TPA (4.31 g, 25.9 mmol) in CH_3_OH (150 mL), 95% H_2_SO_4_ (2.0 mL) was added dropwise at room temperature. The mixture was stirred at reflux for 24 hrs and became a clear solution. The reaction mixture was then cooled to room temperature. DMT was subsequently recrystallized from the reaction mixture and collected after filtration. These white crystals were washed with cold CH_3_OHand dried under vacuum for 4 hrs to afford DMT (4.12 g) with a yield of 82%. ^1^H NMR (400 MHz, CDCl_3_): 8.08 ppm (s, 4H), 3.93 ppm (s, 6H).

To a three-necked round bottom flask equipped with an air condenser, DMT (4.12 g, 21.1 mmol) was added. The flask was evacuated under vacuum and refilled with nitrogen gas for three times. Ethylene glycol (1.20 mL, 21.5 mmol) was added, followed by the addition of titanium isopropoxide (0.06 mL, 0.21 mmol). The mixture was stirred at 160 °C for 1 hour, 200 °C for 1 hour, and 210 °C for 2 hrs under nitrogen. The reaction temperature was further increased to 260 °C, and high vacuum was applied to remove unreacted monomers. The mixture was stirred at 260 °C for 2 hrs and then cooled to room temperature. The resulting PET was dissolved in 1,1,1,3,3,3-hexafluoro-2-propanol (10 mL) and added dropwise to CH_3_OH to remove the catalyst. PET (2.83 g) was collected as white solids after centrifuging and dried under vacuum.

### Simultaneous bioprocess combing *P. putida* Go19 and exogenous PHEs

The simultaneous process experiments were performed using nitrogen-limiting M9 medium (NL-M9) containing M9 Minimal Salts (Sigma-Aldrich, St Louis, MO), 1 g/L NaNH_4_HPO_4_‧4H_2_O, 0.34 g/L thiamine hydrochloride, 2 mM MgSO_4_, and 0.1 mM CaCl_2_. *P. putida* Go19 (accession number NCIMB 41537) was purchased from National Collection of Industrial, Food and Marine Bacteria (NCIMB Aberdeen, Scotland, UK). A single colony of *P. putida* Go19 was inoculated into 20 mL of LB medium and grown overnight at 30 °C/225 rpm. The overnight-grown culture was centrifuged at 4000 g for 5 mins. Cell pellets were then resuspended in 20 mL of NL-M9 medium without any carbon source. The resuspended cells used as inoculum for simultaneous process condition where *P. putida* Go19 and exogenous PHEs were combined to explore the possibility of simultaneous PET depolymerization and fermentation. Subsequently, *P. putida* Go19 was inoculated into 1 mL of NL-M9 medium (to an OD_600_ of 0.2) supplemented with an unpretreated circular pc-PET film (hole-punched from a cookies plastic container; 6 mm in diameter and weigh around 6 mg) absent of any other carbon source in a culture tube. 200 nM of exogenous enzyme (FAST-PETase, wild-type PETase, ThermoPETase, DuraPETase, LCC or ICCM) was added to the 1 mL NL-M9 medium inoculated with *P. putida* Go19 as the simultaneous process condition. 200 nM of enzymes were also respectively added to the 1 mL of NL-M9 medium absent of *P. putida* Go19 as the control condition. All conditions were incubated at 37 ⁰C and 165 RPM for 72 hrs, followed by incubation at 30 ⁰C and 225 RPM for another 24 hrs to maximize PHAs production in *P. putida* Go19. The experiments were all performed in biological triplicates. PET monomers concentration of the NL-M9 medium with or without *P. putida* Go19, mass loss of the pc-PET films, PHAs accumulation and cell density of *P. putida* Go19 were determined after 96 hrs of incubation.

### Analytical method for measuring PET monomers released

HPLC was used to analyze the PET monomers– terephthalic acid (TPA) and Mono-(2-hydroxyethyl)terephthalic (MHET) released from PET depolymerization. The assay samples were filtered with 0.2-μm nylon syringe filters (Wheaton Science, Millville, NJ) prior to running HPLC. Measurement of TPA and MHET was performed using a Dionex UltiMate 3000 (Thermo Fisher Scientific, Waltham, MA) equipped with an Agilent Eclipse Plus C18 column (3.0 × 150 mm, 3.5 μm) with detection wavelength at 260 nm. Column oven was held at 30 °C with 1% acetic acid in water or 1% acetic acid in acetonitrile as mobile phase over the course of the 30-min sequence under the following conditions: 1% to 5% organic (vol/vol) for 5 min, 5% to 100% organic (vol/vol) for 8 min, 100% organic (vol/vol) for 10 min, 100% to 5% organic for 2 min followed by 5% organic for 5 min. The flow rate was fixed at 0.8 mL min^-1^. A standard curve was prepared using commercial TPA with ≥ 98.0% purity or MHET ≥ 98.0% purity (Sigma-Aldrich, St Louis, MO).

### Analytical method for measuring melting temperature (T_m_). of proteins

To evaluate the thermostability of wild-type PETase and its variants, Differential Scanning Calorimetry (DSC) was used to determine their respective T_m_.Purified protein samples were concentrated to 300∼500 µM using Microcon® Centrifugal Filter 10K Devices (Millipore, Billerica, MA). 10 µL of concentrated protein solution (DPBS buffer pH 7.0) was placed in an aluminum Tzero pan and sealed with a Tzero hermetic lid (TA Instruments, DE New Caste, DE). T_m_ of the protein samples was analyzed by a DSC250 (TA Instruments, New Castle, DE) with a RCS90 electric chiller. Two DSC procedures were used depending on the anticipated denaturation temperature of the protein. The first DSC method heated from 40 °C to 90 °C at 10 °C/min, held at 90 °C for two minutes, then cooled from 90 °C to 40 °C at –10 °C/min. The second method heated from 20 °C to 70 °C at 10 °C/min, held at 70 °C for two minutes, cooled from 70 °C to 40 °C at –10 °C/min. The denaturation temperature of the proteins was measured on the heating ramp trace as the midpoint value at half-height.

### Analytical method for determining PET film crystallinity

DSC was used to determine percent crystallinity of the PET films hole-punched from the post-consumer plastics. PET film samples (4–10.5 mg) were placed in aluminum Tzero pans with a Tzero solid sample lid. Samples were run first heated from 40 °C to 300 °C at 5 °C/min, held at 300 °C for one minute, cooled from 300 °C to 30 °C at –5 °C/min, held at 30 °C for one minute in a DSC250 (TA Instruments, New Castle, DE) with a RCS90 electric chiller. The percent crystallinity was determined on the first heating scan using the enthalpies of melting and cold crystallization. The equation used to calculate percent crystallinity within the PET film was the following

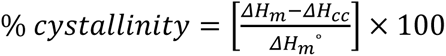

where ΔH_m_ is the enthalpy of melting (J/g), ΔH_cc_ is the enthalpy of cold crystallization (J/g), and ΔH_m_° is the enthalpy of melting for a 100% crystalline PET sample, which is 140.1 J/g^4,27^. ΔH_m_ and ΔH_cc_ were measured by integrating from 90–100 °C to *ca.* 260 °C with a linear baseline. The percent crystallinity was calculated using the TRIOS software package. The glass transition temperatures of the PET films were measured using the second heating scan: 30–300 °C at 10 °C/min, held at 300 °C for one minute, then 300–40 °C at –10 °C/min. The glass transition temperatures for all the PET films were between 80–82 °C.

### Gel Permeation Chromatography (GPC)

GPC measurement was performed at an on an Agilent system with a 1260 Infinity isocratic pump, degasser, and thermostated column chamber held at 30 °C. A mixture of chloroform with 50 ppm amylene and 1,1,1,3,3,3-hexafluoro-2-propanol (2.0 vol%) was used as the mobile phase at 0.5 mL/min. Molecular weights (Mn, Mw) and polydispersity indices (Ð) were determined relative to polystyrene standards.

Sample preparation for GPC

About 8 mg of PET was dissolved in 0.15 mL 1,1,1,3,3,3-hexafluoro-2-propanol. Once PET was completely dissolved, chloroform was added to make a total volume of 2.50 mL. The solution was filtered through a PTFE membrane with a pore size of 0.45 μm. 100 μL of the sample solution was then injected into the GPC system.

### Scanning Electron Microscopy (SEM)

PET films were mounted onto a 3.2 mm SEM stub using carbon tape. Samples were sputter-coated with 8 nm platinum/palladium using a Cressington 208HR Sputter Coater. The metal was sputter coated onto the sample *via* plasma generated with argon present in the chamber. SEM imaging was performed under vacuum using a Zeiss Supra40 Scanning Electron Microscope (SEM). The electron beam intensity was 5 kV.

### Atomic force Microscopy (AFM)

AFM was performed on an Aylum MFP-3D atomic force microscope (Asylum Research, Santa Barbara, CA) in tapping mode. Images were recorded after a surface scan on an area of 50×50 μm. Image analysis including histogram, and surface roughness was performed using Igor Pro.

### Nuclear magnetic resonance (NMR) spectroscopy

^1^H NMR spectroscopy was performed at a 400 MHz Bruker AVANCE NEO spectrometer at room temperature in CDCl_3_ or d_6_-DMSO. Chemical shift was referenced to the residual solvent signal (^1^H NMR: 7.26 ppm in CDCl_3_, 2.50 ppm in d_6_-DMSO, respectively).

### Microscopic observation of *P. putida* Go19 and PHAs

PHAs formation in *P. putida* Go19 from the simultaneous experiment was microscopically observed by using an upright laser scanning confocal microscope (LSM-510, Zeiss) under reflectance using a 100X, 1.3 NA oil-dipping objective (Zeiss), according to the protocol described by Franden et al.^28^. 0.5 mL sample was taken from the simultaneous process experiment of the *P. putida* Go19 culture (with 200 nM of FAST-PETase) that grew on NL-M9 medium supplemented with a circular pc-PET film (cookie container) as the sole carbon source. Culture was centrifuged at 4,000 g for 3 min, followed by washing twice with DPBS. Cell pellets were then stained with 1 mL of 10 μg/mL Nile Red (Thermo Fisher Scientific, Waltham, MA, USA) and kept in the dark for 30 min at room temperature. Subsequently, the stained cells were pelleted again by centrifugation at 4,000 g for 3 min, followed by washing twice with DPBS. Finally, the stained cells were resuspended in 200 μL of DPBS. 1 ul of the stained cell solution was dipped on a coverslip and pressed with a small coverslip. The samples were illuminated with a 561 nm laser and reflected light was collected by setting the Meta detector channel between 570-670 nm wavelength.

## Supplementary method 2

Sequences used in this study

### Nucleotide sequence of the signal peptide—SPpstu (21 amino acids) from maltotetraose-forming amylase of *Pseudomonas stutzeri* MO-19^25^

atgagccacatcctgcgagccgccgtattggcggcgatgctgttgccgttgccgtccatggcc

### Wild-type PETase

#### Nucleotide sequence of wild-type PETase without its original signal peptide

cagaccaatccgtatgcgcgcggtccgaatccgacagccgccagtttggaagcgagcgctggtccattcaccgttcgctcctttaccgtgagtagaccgagcggttatggcgctggcaccgtttactatccaacaaatgctgggggtaccgtgggcgccatagccatagttcccgggtatacggcacggcagtcatcaattaaatggtggggaccgcgtctggcatcccacggtttcgtagtaattacaattgacacaaattccacgttagaccagccatcaagtcggagttcgcaacaaatggccgcgctgcgccaggtggcgtcgttaaacggtacaagtagcagcccgatttacggaaaggtcgataccgctcgtatgggtgttatggggtggagtatgggaggtggaggctccctgatctctgctgctaacaacccttcgctgaaagcagcggcgcctcaagcaccatgggattcttcgacaaattttagttctgtaactgtgcccacgctgatcttcgcatgtgaaaacgatagtatagccccggtcaactcttcagcacttcctatctatgattctatgtcacgcaacgctaagcagtttctcgaaattaatggtggctcacattcctgtgcgaatagcggcaattctaaccaagcattaatcggaaaaaaaggcgttgcatggatgaaacgttttatggacaatgatactaggtattctacttttgcctgcgagaacccgaatagcaccagagtgtctgattttcgtacagcgaattgcagcctcgagcaccaccaccaccaccac

#### Expressed amino acid sequence of wild-type PETase by *P. putida* GO16/KT2440

QTNPYARGPNPTAASLEASAGPFTVRSFTVSRPSGYGAGTVYYPTNAGGTVGAIAIVPGY TARQSSIKWWGPRLASHGFVVITIDTNSTLDQPSSRSSQQMAALRQVASLNGTSSSPIYGK VDTARMGVMGWSMGGGGSLISAANNPSLKAAAPQAPWDSSTNFSSVTVPTLIFACENDSI APVNSSALPIYDSMSRNAKQFLEINGGSHSCANSGNSNQALIGKKGVAWMKRFMDNDTR YSTFACENPNSTRVSDFRTANCSLEHHHHHH

### ThermoPETase

#### Nucleotide sequence of ThermoPETase without its original signal peptide sequence

cagaccaatccgtatgcgcgcggtccgaatccgacagccgccagtttggaagcgagcgctggtccattcaccgttcgctcctttaccgtgagtagaccgagcggttatggcgctggcaccgtttactatccaacaaatgctgggggtaccgtgggcgccatagccatagttcccgggtatacggcacggcagtcatcaattaaatggtggggaccgcgtctggcatcccacggtttcgtagtaattacaattgacacaaattccacgttagaccagccagaaagtcggagttcgcaacaaatggccgcgctgcgccaggtggcgtcgttaaacggtacaagtagcagcccgatttacggaaaggtcgataccgctcgtatgggtgttatggggtggagtatgggaggtggaggctccctgatctctgctgctaacaacccttcgctgaaagcagcggcgcctcaagcaccatggcactcttcgacaaattttagttctgtaactgtgcccacgctgatcttcgcatgtgaaaacgatagtatagccccggtcaactcttcagcacttcctatctatgattctatgtcacgcaacgctaagcagtttctcgaaattaatggtggctcacattcctgtgcgaatagcggcaattctaaccaagcattaatcggaaaaaaaggcgttgcatggatgaaacgttttatggacaatgatactaggtattctacttttgcctgcgagaacccgaatagcaccgcagtgtctgattttcgtacagcgaattgcagcctcgagcaccaccaccaccaccac

#### Expressed amino acid sequence of ThermoPETase by *P. putida* KT2440

QTNPYARGPNPTAASLEASAGPFTVRSFTVSRPSGYGAGTVYYPTNAGGTVGAIAIVPGY TARQSSIKWWGPRLASHGFVVITIDTNSTLDQPESRSSQQMAALRQVASLNGTSSSPIYGK VDTARMGVMGWSMGGGGSLISAANNPSLKAAAPQAPWHSSTNFSSVTVPTLIFACENDSI APVNSSALPIYDSMSRNAKQFLEINGGSHSCANSGNSNQALIGKKGVAWMKRFMDNDTR YSTFACENPNSTAVSDFRTANCSLEHHHHHH

### DuraPETase

#### Nucleotide sequence of DuraPETase without its original signal peptide sequence

cagaccaatccgtatgcgcgcggtccgaatccgacagccgccagtttggaagcgagcgctggtccattcaccgttcgctcctttaccgtgagtagaccgagcggttatggcgctggcaccgtttactatccaacaaatgctgggggtaccgtgggcgccatagccatagttcccgggtatacggcacggcagtcatcaattaaatggtggggaccgcgtctggcatcccacggtttcgtagtaattacaattgacacaaattccacgtttgactatccatcaagtcggagttcgcaacaaatggccgcgctgcgccaggtggcgtcgttaaacggtgacagtagcagcccgatttacggaaaggtcgataccgctcgtatgggtgttatggggcatagtatgggaggtggagcatccctgcgatctgctgctaacaacccttcgctgaaagcagcgattcctcaagcaccatgggattctcaaacaaattttagttctgtaactgtgcccacgctgatcttcgcatgtgaaaacgatagtatagccccggtcaactctcatgcacttcctatctatgattctatgtcacgcaacgctaagcagtttctcgaaattaatggtggctcacattcctgtgcgaatagcggcaattctaaccaagcattaatcggaaaaaaaggcgttgcatggatgaaacgttttatggacaatgatactaggtattctacttttgcctgcgagaacccgaatagcaccgcagtgtctgattttcgtacagcgaattgcagcctcgagcaccaccaccaccaccac

#### Expressed amino acid sequence of DuraPETase by *P. putida* KT2440

QTNPYARGPNPTAASLEASAGPFTVRSFTVSRPSGYGAGTVYYPTNAGGTVGAIAIVPGY TARQSSIKWWGPRLASHGFVVITIDTNSTFDYPSSRSSQQMAALRQVASLNGDSSSPIYGK VDTARMGVMGHSMGGGASLRSAANNPSLKAAIPQAPWDSQTNFSSVTVPTLIFACENDSI APVNSHALPIYDSMSRNAKQFLEINGGSHSCANSGNSNQALIGKKGVAWMKRFMDNDTR YSTFACENPNSTAVSDFRTANCSLEHHHHHH

### LCC

#### Nucleotide sequence of LCC without its original signal peptide sequence

agcaacccgtaccagcgtggcccgaatccgacccgcagcgcactgaccgcagatggcccgtttagcgtggcaacctacaccgtctcacgcctgtcagtctcgggttttggcggtggcgtgatttattacccgaccggcacgtctctgacgttcggtggcatcgcgatgagtccgggttataccgcagatgctagctctctggcatggctgggtcgtcgcctggcttcccatggctttgtggttctggtgattaacacgaattcacgtttcgattatccggacagccgcgcctctcagctgagtgccgccctgaactacctgcgtaccagttccccgagcgccgttcgcgcacgtctggatgcaaatcgtctggcggttgccggtcattctatgggtggcggtggcaccctgcgtattgcagaacaaaacccgagcctgaaagcggctgtcccgctgaccccgtggcacaccgataaaacgtttaataccagtgtcccggtgctgattgttggcgcagaagctgacaccgtggcgccggtttcgcagcatgccatcccgttttatcaaaacctgccgagcaccacgccgaaagtttacgtcgaactggataacgcatcgcacttcgctccgaatagcaacaatgcggccatttccgtttatacgatctcatggatgaaactgtgggtcgataatgacacccgttaccgccagttcctgtgtaatgtgaacgacccggctctgtccgacttccgcaccaataatcgccactgccaactcgagcaccaccaccaccaccac

#### Expressed amino acid sequence of LCC by *E. coli*

QTNPYARGPNPTAASLEASAGPFTVRSFTVSRPSGYGAGTVYYPTNAGGTVGAIAIVPGY TARQSSIKWWGPRLASHGFVVITIDTNSTFDYPSSRSSQQMAALRQVASLNGDSSSPIYGK VDTARMGVMGHSMGGGASLRSAANNPSLKAAIPQAPWDSQTNFSSVTVPTLIFACENDSI APVNSHALPIYDSMSRNAKQFLEINGGSHSCANSGNSNQALIGKKGVAWMKRFMDNDTR YSTFACENPNSTAVSDFRTANCSLEHHHHHH

### ICCM

#### Nucleotide sequence of ICCM without its original signal peptide sequence

atgagcaacccgtaccagcgtggcccgaatccgacccgcagcgcactgaccgcagatggcccgtttagcgtggcaacctacaccgtctcac gcctgtcagtctcgggttttggcggtggcgtgatttattacccgaccggcacgtctctgacgttcggtggcatcgcgatgagtccgggttatacc gcagatgctagctctctggcatggctgggtcgtcgcctggcttcccatggctttgtggttctggtgattaacacgaattcacgtttcgattatccgg acagccgcgcctctcagctgagtgccgccctgaactacctgcgtaccagttccccgagcgccgttcgcgcacgtctggatgcaaatcgtctg gcggttgccggtcattctatgggtggcggtggcaccctgcgtattgcagaacaaaacccgagcctgaaagcggctgtcccgctgaccccgtg gcacaccgataaaacgtttaataccagtgtcccggtgctgattgttggcgcagaagctgacaccgtggcgccggtttcgcagcatgccatccc gttttatcaaaacctgccgagcaccacgccgaaagtttacgtcgaactgtgcaacgcatcgcacattgctccgatgagcaacaatgcggccatt tccgtttatacgatctcatggatgaaactgtgggtcgataatgacacccgttaccgccagttcctgtgtaatgtgaacgacccggctctgtgcgac ttccgcaccaataatcgccactgccaactcgagcaccaccaccaccaccac

#### Expressed amino acid sequence of LCC by *E. coli*

MSNPYQRGPNPTRSALTADGPFSVATYTVSRLSVSGFGGGVIYYPTGTSLTFGGIAMSPGY TADASSLAWLGRRLASHGFVVLVINTNSRFDYPDSRASQLSAALNYLRTSSPSAVRARLD ANRLAVAGHSMGGGGTLRIAEQNPSLKAAVPLTPWHTDKTFNTSVPVLIVGAEADTVAP VSQHAIPFYQNLPSTTPKVYVELCNASHIAPMSNNAAISVYTISWMKLWVDNDTRYRQFL CNVNDPALCDFRTNNRHCQLEHHHHHH

## Legends for Supplementary Information Figures

**Supplementary Information Fig. 1 | Schematic diagram of Mutcompute. a.** Creating a microenvironment: MutCompute begins by centering itself on the alpha carbon of a particular residue in the protein and filters all peptide atoms within a 20 angstrom cube (the orientation of the cube is normalized with respect to the protein backbone). In the filtering process, we create an artificial, self-supervised label by excluding all atoms that belong the center residue. **b.** Encoding the microenvironment: The filtered atoms are then encoded into a 7-channel voxelated representation with a voxel resolution of 1A^3^. **c.** Running MutCompute on a Microenvironment: The 7-channel voxelated representation of a microenvironment is then passed to the CNN model, MutCompute. The model can be broken into 2 parts: Feature extraction and classification. The feature extraction portion consist of convolutional and max pooling layers and is then flattened into a 1D-vector before being passed to the classification layers of the model. The output is a probability mass function of the likelihood each of the 20 amino acids was the amino acid in the center of the microenvironment. We do this process for every residue in the protein to identify residues for mutagenesis.

**Supplementary Information Fig. 2 | Disfavored PETase residues flagged by MutCompute from the wild-type and ThermoPETase crystal structures.** MutCompute outputs a probability distribution that describes the likelihood of each of the 20 canonical amino acids to be the wild-type amino acid for the surrounding chemical environment. A disfavored residue is defined as a residue where the amino acid with the highest predicted probability is not the wild-type amino acid. Here, a 30% wild-type probability cutoff was used to down select disfavored residues.

**Supplementary Information Fig. 3a | Predictions (based on wild-type PETase) ranked by fold change in the probabilities between the predicted and the wild-type amino acid.** Fold change predictions are provided as a means of down-selecting potential mutations.

**Supplementary Information Fig. 3b | TOP 10 ranked predictions (based on wild-type PETase).** The top 10 mutations predicted for the wild-type PETase scaffold are presented.

**Supplementary Information Fig. 3c | Predictions (based on ThermoPETase) ranked by fold change in the probabilities between the predicted and the wild-type amino acid.** Fold change predictions are provided as a means of down-selecting potential mutations.

**Supplementary Information Fig. 3d | TOP 10 ranked predictions (based on ThermoPETase).** The top 10 mutations predicted for the ThermoPETase scaffold are presented.

**Supplementary Information Fig. 4 | Selecting mutations based on experimental catalytic activity measurements.** A scheme for selecting mutations based on experimental evidence is provided.

**Supplementary Information Fig. 5 | Thermostability of the PETase variants incorporating the mutations predicted by Mutcompute and their respective scaffolds—wild-type PETase (WT), ThermoPETase (Thermo), DuraPETase (Dura).** The melting temperature of each enzyme was determined by DSC. All measurement were conducted in triplicate (n=3).

**Supplementary Information Fig. 6 | Protein yield of the PETase variants incorporating the mutations predicted by Mutcompute and their respective scaffolds—wild-type PETase (WT), ThermoPETase (Thermo), DuraPETase (Dura).** Protein yields from *P. putida* purification experiments indicate improved yields from mutant enzymes.

**Supplementary Information Fig. 7 | The PET-hydrolytic activity of FAST-PETase outperformed various PHEs at mild temperatures and modest pH.** Comparison of PET-hydrolytic activity of FAST-PETase, wild-type PETase (WT), ThermoPETase (Thermo), DuraPETase (Dura), LCC and ICCM across a range of pH (6.5 – 8.0) at reaction temperatures of 30 ⁰C (**a**.) and 40 ⁰C (**b**.). PET-hydrolytic activity was evaluated by measuring the amount of PET monomers (the sum of TPA and MHET) released from hydrolyzing gf-PET film by the tested enzymes after 96 hrs of reaction time. All measurement were conducted in triplicate (n=3).

**Supplementary Information Fig. 8 | Mass, crystallinity %, molecular weights (*M*_n_, *M*_w_), polydispersity indices (Ð) and time for complete degradation of various pc-PET films by FAST-PETase.** The circular pc-PET films (6 mm in diameter) were hole-punched from 51 different post-consumer plastic products used in the packaging of food, beverages, medications, office supplies, household goods and cosmetics available at local grocery store chains (Walmart, Costco, and HEB). The pc-PET films were hydrolysed by serial treatment with FAST-PETase at 50 ⁰C until the films were completed degraded. The enzyme solution (200 nM of FAST-PETase in 100mM KH_2_PO_4_-NaOH (pH 8.0) buffer) was replenished every 24 hours. The crystallinity % of the intact pc-PET films was determined by DSC. The initial mass of the films was determined gravimetrically by a digital scale. Both DSC and gravimetric measurements were conducted in triplicate. Means ± s.d. (n=3) are shown.

**Supplementary Information Fig. 9 | Scatterplot of degradation rate versus (a.) initial mass or (b.) crystallinity % or (c.) weight average molecular weight (Mw) or (d.) number average molecular weight (Mn) or (e.) polydispersity indices of the hole-punched films from 51 different post-consumer plastic products.** Degradation rate was not found to be dependent on any one metric of these various plastics.

**Supplementary Information Fig. 10 | Time-course of crystallinity % of the degraded pc-PET film.** The hole-punched PET films from a bean cake PET container were treated with FAST-PETase for 0 hr, 4hr, 8 hr,12 hrs 16 hr in 100 mM KH_2_PO_4_-NaOH(pH 8.0) buffer at 50 ⁰C. Crystallinity % of the films was determined by DSC. All measurement were conducted in duplicate (n=2).

**Supplementary Information Fig. 11 | Scanning electron microscopic analysis of the pc-PET films.** The hole-punched PET films from a bean cake PET container were treated with FAST-PETase for 0 hr, 8 hr, 16 hr in 100 mM KH_2_PO_4_-NaOH (pH 8.0) buffer at 50 ⁰C.

**Supplementary Information Fig. 12 | The surface roughness of the pc-PET films determined by atomic force microscopy.** The hole-punched PET films from a bean cake PET container were treated with FAST-PETase for 4 hr, 8 hr, 12 hr, 16 hr and 20 hr in 100mM KH_2_PO_4_-NaOH (pH 8.0) buffer at 50 ⁰C. The time-course profile of the surface roughness indicated that longer exposure times with FAST-PETase resulted in higher degree of surface roughness on the pc-PET films. RMS represents root mean square.

**Supplementary Information Fig. 13 | Time-course of PET-hydrolytic activity of LCC and ICCM at reaction temperatures of 55 ⁰C, 60 ⁰C, 65 ⁰C, and 72 ⁰C.** PET-hydrolytic activity was evaluated by measuring the amount of PET monomers (the sum of TPA and MHET) released from hydrolyzing the pc-PET (Bean cake plastic container) film by the tested PHEs at various time points. 100 mM KH_2_PO_4_-NaOH(pH 8.0) buffer was used for all reactions shown in this figure. All measurement were conducted in triplicate (n=3).

**Supplementary Information Fig. 14 | A closed-loop PET recycling process**. Demonstration of a closed-loop process for enzymatically degrading and then regenerating PET in the course of several days.

**SupplementaryInformationFig. 15 | a. ^1^H NMR (400 MHz, *d*_6_-DMSO)spectraof TPA recovered from degraded PET solutions.** The peak at 8.029 ppm corresponds to the hydrogen nuclei of the benzenering.**b. ^1^H NMR (400 MHz, CDCl_3_) spectra of DMT synthesized from TPA.** The peak at 8.081 ppm corresponds to the hydrogen nuclei of the benzene ring. The peak at 3.93 ppm corresponds to the hydrogen nuclei of the methyl group.

**Supplementary Information Fig. 16 | DSC trace of PET regenerated from the degraded solutions.** The crystallinity of this regenerated PET is 58.46%. The melting onset is 243.6 °C. The melting peak temperature is 258.4 °C. The glass transition temperature is 84.3 °C.

**Supplementary Information Fig. 17 | X-ray crystal structure of FAST-PETase. a.** Overall crystal structure of FAST-PETase. Catalytic triads (S160, D206, H237) are shown in blue sticks. Mutations originating from ThermoPETase (S121E, D186H, R280A) are shown in pink sticks, and novel mutationspredictedbythe neural network are shown in green-yellow sticks. **b-c.** 2F_o_-F_c_ map (contoured at 1.5σ) shown as grey mesh superimposed on the stick models of novel mutation sites (b.) R224Q, (c.) N233K.

**Supplementary Information Fig. 18 | Statistics of the crystal structural determination of FAST-PETase.** Information about the obtained crystal structure is provided.

**Supplementary Information Fig. 19 | Stages of degradation of pc-PET films by FAST-PETase.** a. The transparent pc-PET film (6 mm in diameter) was completely degraded (only cutting edges of the film remained) after 24 hrs of a single treatment with FAST-PETase at 50 ⁰C. b. The colored pc-PET film (6 mm in diameter) was completely degraded (only cutting edges of the film and some colorants remained) after six day of serial treatment with FAST-PETase at 50 ⁰C. Enzyme (200 nM) treatment was performed with 100 mM KH_2_PO_4_-NaOH (pH 8.0) buffer.

