## Supplementary Information for "Deep learning redesign of PETase for practical PET degrading applications"

#### Affiliations

#### Contents:

Supplementary Information Figures 1-19

### Supplementary Information Figures

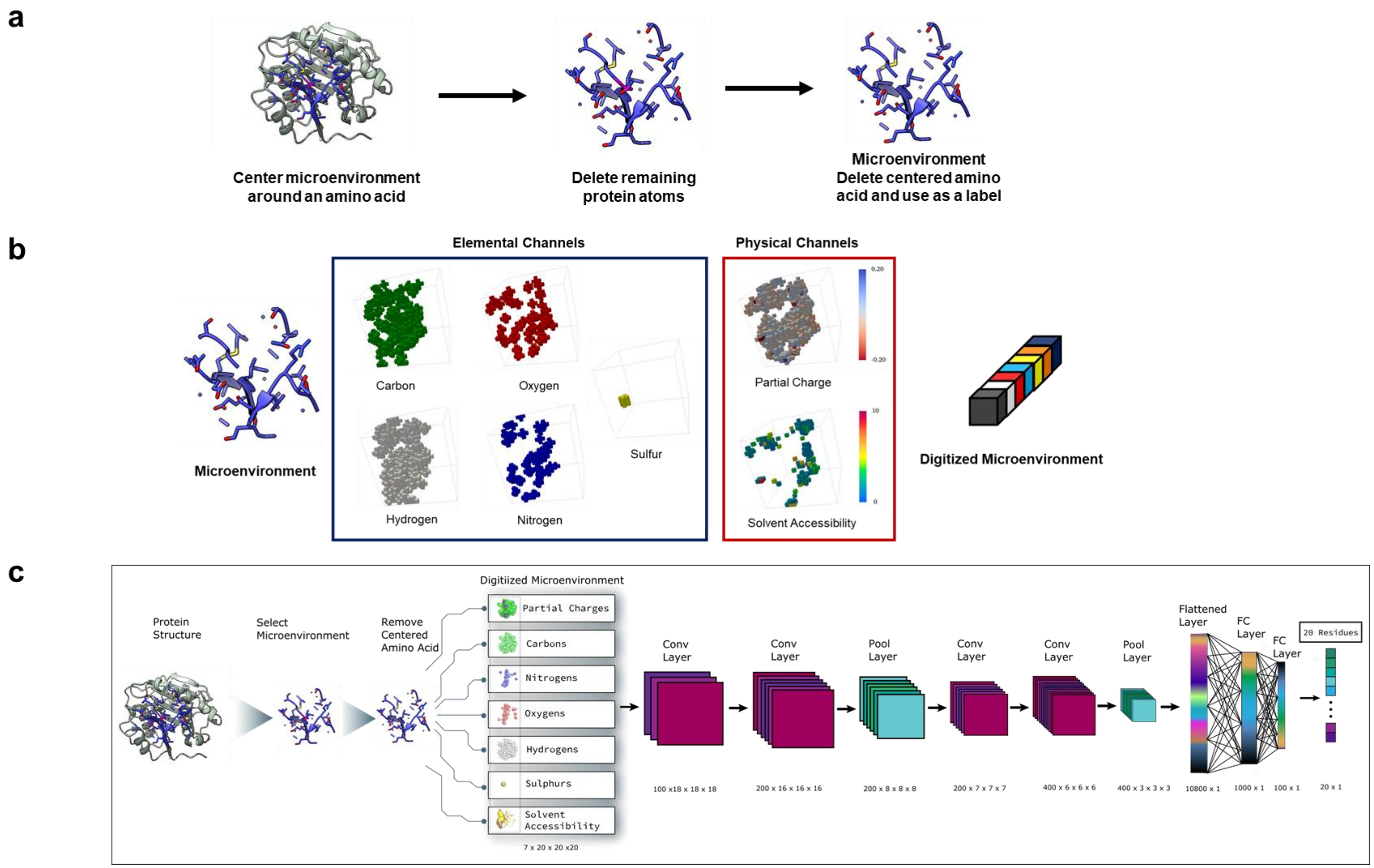

**Supplementary Information Fig. 1 | Schematic diagram of Mutcompute. a.** Creating a microenvironment: MutCompute begins by centering itself on the alpha carbon of a particular residue in the protein and filters all peptide atoms within a 20 angstrom cube (the orientation of the cube is normalized with respect to the protein backbone). In the filtering process, we create an artificial, self-supervised label by excluding all atoms that belong the center residue. **b.** Encoding the microenvironment: The filtered atoms are then encoded into a 7-channel voxelated representation with a voxel resolution of $1A^3$. **c.** Running MutCompute on a Microenvironment: The 7-channel voxelated representation of a microenvironment is then passed to the CNN model, MutCompute. The model can be broken into 2 parts: Feature extraction and classification. The feature extraction portion consist of convolutional and max pooling layers and is then flattened into a 1D-vector before being passed to the classification layers of the model. The output is a probability mass function of the likelihood each of the 20 amino acids was the amino acid in the center of the microenvironment. We do this process for every residue in the protein to identify residues for mutagenesis.

| | | | | Probability distribution across all 20 amino acids | | | | | | | | | | | | | | | | | | | |
|---|---|---|---|---|---|---|---|---|---|---|---|---|---|---|---|---|---|---|---|---|---|---|---|
| **pdb_id** | **pos** | **wtAA** | **wt_prob** | **ALA** | **ARG** | **ASN** | **ASP** | **CYS** | **GLN** | **GLU** | **GLY** | **HIS** | **ILE** | **LEU** | **LYS** | **MET** | **PHE** | **PRO** | **SER** | **THR** | **TRP** | **TYR** | **VAL** |
| 6ij6 | 233 | ASN | 0.09% | 1.E-05 | 1.E-02 | 9.E-04 | 2.E-04 | 2.E-05 | 5.E-03 | 4.E-03 | 3.E-07 | 6.E-04 | 1.E-03 | 5.E-04 | 1.E+00 | 4.E-04 | 2.E-04 | 1.E-06 | 8.E-05 | 3.E-04 | 2.E-05 | 1.E-04 | 9.E-04 |
| 6ij6 | 91 | GLN | 0.09% | 9.E-06 | 6.E-05 | 6.E-05 | 9.E-07 | 3.E-05 | 9.E-04 | 3.E-05 | 6.E-08 | 2.E-07 | 1.E+00 | 4.E-05 | 2.E-04 | 1.E-04 | 3.E-09 | 2.E-07 | 2.E-05 | 6.E-03 | 1.E-08 | 1.E-08 | 1.E-02 |
| 6ij6 | 225 | ASN | 0.32% | 3.E-04 | 9.E-06 | 3.E-03 | 7.E-07 | 1.E+00 | 2.E-05 | 9.E-07 | 4.E-07 | 4.E-07 | 1.E-05 | 1.E-04 | 3.E-07 | 1.E-04 | 3.E-08 | 2.E-07 | 2.E-04 | 2.E-02 | 4.E-09 | 2.E-09 | 1.E-03 |
| 5xjh | 140 | THR | 0.41% | 7.E-05 | 2.E-06 | 1.E-02 | 1.E+00 | 2.E-03 | 3.E-05 | 6.E-04 | 5.E-07 | 2.E-06 | 5.E-07 | 1.E-05 | 1.E-05 | 3.E-06 | 1.E-07 | 1.E-07 | 1.E-02 | 4.E-03 | 9.E-08 | 2.E-07 | 2.E-05 |
| 6ij6 | 61 | SER | 0.54% | 4.E-04 | 9.E-04 | 3.E-02 | 2.E-01 | 2.E-03 | 1.E-02 | 9.E-02 | 9.E-06 | 9.E-03 | 2.E-02 | 2.E-02 | 1.E-03 | 1.E-03 | 1.E-03 | 2.E-05 | 5.E-03 | 3.E-01 | 2.E-04 | 2.E-03 | 3.E-01 |
| 5xjh | 121 | SER | 0.70% | 2.E-03 | 2.E-02 | 3.E-02 | 4.E-02 | 2.E-03 | 1.E-01 | 5.E-01 | 3.E-05 | 7.E-03 | 5.E-04 | 2.E-03 | 2.E-01 | 2.E-03 | 2.E-04 | 1.E-03 | 7.E-03 | 6.E-04 | 2.E-05 | 1.E-04 | 7.E-04 |
| 5xjh | 58 | SER | 0.81% | 4.E-03 | 2.E-02 | 3.E-02 | 1.E-01 | 1.E-03 | 6.E-02 | 5.E-01 | 9.E-05 | 3.E-03 | 7.E-03 | 8.E-02 | 1.E-01 | 2.E-03 | 5.E-04 | 4.E-05 | 8.E-03 | 4.E-03 | 1.E-04 | 3.E-04 | 7.E-03 |
| 5xjh | 95 | LYS | 1.06% | 2.E-04 | 8.E-04 | 1.E-02 | 2.E-02 | 2.E-03 | 2.E-01 | 6.E-01 | 2.E-07 | 2.E-05 | 4.E-02 | 1.E-03 | 1.E-02 | 2.E-03 | 9.E-07 | 5.E-07 | 9.E-03 | 4.E-03 | 4.E-07 | 9.E-07 | 1.E-01 |
| 5xjh | 279 | THR | 1.98% | 9.E-03 | 2.E-02 | 1.E-01 | 5.E-01 | 2.E-03 | 3.E-02 | 2.E-01 | 2.E-04 | 9.E-03 | 5.E-03 | 3.E-03 | 3.E-02 | 2.E-03 | 1.E-03 | 1.E-04 | 8.E-02 | 2.E-02 | 2.E-04 | 8.E-04 | 9.E-03 |
| 5xjh | 263 | ASP | 2.25% | 7.E-05 | 2.E-05 | 5.E-02 | 2.E-02 | 4.E-04 | 5.E-02 | 8.E-03 | 9.E-07 | 9.E-01 | 7.E-07 | 2.E-05 | 5.E-04 | 6.E-03 | 2.E-04 | 2.E-07 | 9.E-05 | 1.E-04 | 2.E-06 | 5.E-04 | 6.E-07 |
| 6ij6 | 58 | SER | 2.30% | 8.E-01 | 1.E-02 | 6.E-03 | 3.E-02 | 4.E-04 | 8.E-03 | 1.E-01 | 1.E-03 | 4.E-04 | 2.E-04 | 6.E-03 | 2.E-02 | 6.E-04 | 1.E-04 | 5.E-05 | 2.E-02 | 4.E-03 | 4.E-05 | 2.E-04 | 3.E-03 |
| 6ij6 | 53 | ARG | 2.82% | 1.E-04 | 3.E-02 | 1.E-02 | 4.E-04 | 7.E-05 | 3.E-01 | 7.E-02 | 1.E-06 | 3.E-05 | 3.E-04 | 7.E-03 | 6.E-01 | 8.E-03 | 5.E-06 | 1.E-06 | 2.E-04 | 1.E-04 | 5.E-06 | 6.E-06 | 2.E-04 |
| 6ij6 | 34 | ARG | 2.99% | 4.E-07 | 3.E-02 | 2.E-02 | 2.E-04 | 8.E-05 | 3.E-03 | 5.E-04 | 3.E-07 | 1.E-01 | 3.E-02 | 8.E-01 | 2.E-02 | 4.E-03 | 6.E-04 | 6.E-07 | 2.E-05 | 1.E-03 | 4.E-05 | 9.E-05 | 5.E-04 |
| 6ij6 | 208 | ILE | 3.02% | 4.E-04 | 4.E-03 | 4.E-04 | 3.E-04 | 3.E-04 | 6.E-04 | 8.E-04 | 3.E-06 | 7.E-05 | 3.E-02 | 5.E-04 | 1.E-03 | 5.E-04 | 3.E-05 | 1.E-05 | 6.E-04 | 7.E-02 | 1.E-05 | 1.E-05 | 9.E-01 |
| 6ij6 | 59 | ARG | 3.25% | 2.E-02 | 3.E-02 | 6.E-02 | 5.E-02 | 6.E-03 | 6.E-02 | 1.E-01 | 3.E-04 | 3.E-03 | 4.E-03 | 1.E-03 | 9.E-02 | 3.E-03 | 4.E-04 | 2.E-04 | 2.E-01 | 3.E-01 | 5.E-05 | 5.E-04 | 6.E-02 |
| 6ij6 | 224 | ARG | 3.52% | 1.E-05 | 4.E-02 | 1.E-02 | 3.E-04 | 1.E-05 | 4.E-01 | 1.E-02 | 4.E-07 | 3.E-03 | 6.E-06 | 2.E-01 | 4.E-01 | 1.E-02 | 1.E-03 | 5.E-07 | 8.E-06 | 3.E-06 | 2.E-03 | 1.E-03 | 5.E-07 |
| 5xjh | 262 | MET | 3.74% | 5.E-06 | 8.E-06 | 1.E-02 | 9.E-05 | 2.E-03 | 3.E-04 | 9.E-05 | 5.E-07 | 3.E-05 | 2.E-01 | 7.E-01 | 5.E-05 | 4.E-02 | 3.E-06 | 8.E-07 | 7.E-06 | 6.E-04 | 3.E-07 | 5.E-07 | 5.E-02 |
| 6ij6 | 136 | SER | 4.15% | 2.E-02 | 3.E-05 | 7.E-03 | 1.E-01 | 2.E-02 | 5.E-03 | 4.E-02 | 7.E-05 | 2.E-03 | 1.E-04 | 6.E-04 | 2.E-04 | 5.E-04 | 4.E-04 | 1.E-05 | 4.E-02 | 7.E-01 | 2.E-05 | 2.E-04 | 9.E-02 |
| 5xjh | 233 | ASN | 4.51% | 6.E-04 | 3.E-03 | 5.E-02 | 7.E-03 | 5.E-04 | 2.E-02 | 1.E-02 | 1.E-06 | 7.E-02 | 2.E-05 | 3.E-04 | 8.E-01 | 4.E-04 | 1.E-03 | 7.E-07 | 2.E-03 | 1.E-03 | 1.E-05 | 2.E-03 | 6.E-05 |
| 6ij6 | 186 | HIS | 4.75% | 1.E-05 | 2.E-06 | 2.E-01 | 8.E-01 | 3.E-03 | 2.E-04 | 1.E-03 | 4.E-08 | 5.E-02 | 7.E-07 | 1.E-05 | 2.E-05 | 4.E-05 | 5.E-04 | 2.E-07 | 7.E-04 | 3.E-05 | 1.E-04 | 3.E-04 | 9.E-06 |
| 5xjh | 225 | ASN | 4.83% | 2.E-03 | 2.E-04 | 5.E-02 | 3.E-06 | 6.E-01 | 2.E-04 | 5.E-06 | 1.E-06 | 9.E-06 | 8.E-06 | 4.E-04 | 3.E-06 | 3.E-04 | 7.E-08 | 1.E-06 | 4.E-02 | 3.E-01 | 2.E-08 | 2.E-08 | 2.E-04 |
| 5xjh | 59 | ARG | 5.74% | 7.E-03 | 6.E-02 | 2.E-01 | 6.E-02 | 3.E-03 | 1.E-01 | 2.E-01 | 2.E-04 | 2.E-02 | 3.E-03 | 3.E-03 | 9.E-02 | 6.E-03 | 3.E-03 | 1.E-04 | 5.E-02 | 2.E-01 | 2.E-04 | 3.E-03 | 3.E-02 |
| 5xjh | 212 | ASN | 6.08% | 3.E-01 | 1.E-02 | 6.E-02 | 3.E-01 | 2.E-03 | 6.E-03 | 2.E-02 | 7.E-04 | 4.E-04 | 2.E-04 | 3.E-03 | 3.E-02 | 9.E-04 | 1.E-04 | 3.E-05 | 3.E-01 | 7.E-03 | 2.E-05 | 1.E-04 | 9.E-04 |
| 6ij6 | 37 | ASN | 6.92% | 4.E-05 | 5.E-05 | 7.E-02 | 9.E-01 | 1.E-04 | 1.E-05 | 2.E-04 | 5.E-07 | 9.E-05 | 5.E-06 | 3.E-06 | 2.E-04 | 5.E-06 | 2.E-06 | 8.E-07 | 5.E-04 | 6.E-05 | 2.E-07 | 2.E-06 | 2.E-05 |
| 6ij6 | 33 | MET | 7.37% | 3.E-05 | 4.E-03 | 7.E-03 | 4.E-03 | 4.E-04 | 4.E-01 | 4.E-01 | 4.E-06 | 4.E-02 | 6.E-03 | 7.E-02 | 6.E-02 | 7.E-02 | 4.E-04 | 4.E-05 | 6.E-04 | 3.E-03 | 2.E-05 | 5.E-04 | 1.E-03 |

| | | | | | | | | | | | | | | | | | | | | | | | |
|---|---|---|---|---|---|---|---|---|---|---|---|---|---|---|---|---|---|---|---|---|---|---|---|
| 6ij6 | 279 | THR | 7.58% | 7.E-03 | 1.E-02 | 4.E-02 | 2.E-01 | 2.E-03 | 2.E-02 | 3.E-01 | 6.E-05 | 1.E-02 | 1.E-02 | 3.E-03 | 8.E-02 | 3.E-03 | 2.E-03 | 9.E-05 | 1.E-01 | 8.E-02 | 2.E-04 | 1.E-03 | 3.E-02 |
| 5xjh | 119 | GLN | 7.67% | 1.E-05 | 5.E-03 | 1.E-03 | 3.E-04 | 1.E-04 | 8.E-02 | 7.E-03 | 2.E-04 | 7.E-03 | 3.E-05 | 1.E-01 | 1.E-01 | 5.E-01 | 1.E-01 | 1.E-06 | 2.E-05 | 5.E-05 | 3.E-03 | 3.E-02 | 8.E-06 |
| 5xjh | 125 | SER | 8.20% | 4.E-02 | 4.E-03 | 6.E-03 | 3.E-02 | 2.E-02 | 2.E-02 | 4.E-02 | 5.E-05 | 2.E-04 | 7.E-03 | 2.E-03 | 1.E-02 | 2.E-03 | 7.E-05 | 5.E-05 | 8.E-02 | 2.E-01 | 3.E-05 | 8.E-05 | 5.E-01 |
| 6ij6 | 212 | ASN | 8.26% | 2.E-01 | 3.E-02 | 8.E-02 | 1.E-01 | 4.E-03 | 3.E-02 | 1.E-01 | 3.E-04 | 2.E-02 | 9.E-03 | 3.E-03 | 7.E-02 | 2.E-03 | 3.E-03 | 5.E-05 | 2.E-01 | 5.E-03 | 2.E-04 | 2.E-03 | 1.E-02 |
| 6ij6 | 87 | TYR | 8.43% | 8.E-08 | 5.E-01 | 7.E-06 | 1.E-06 | 2.E-06 | 1.E-03 | 1.E-04 | 4.E-08 | 1.E-02 | 1.E-05 | 3.E-03 | 4.E-01 | 4.E-03 | 2.E-02 | 5.E-08 | 2.E-07 | 1.E-07 | 2.E-03 | 8.E-02 | 2.E-07 |
| 5xjh | 82 | ALA | 8.52% | 9.E-02 | 1.E-07 | 1.E-05 | 2.E-06 | 5.E-03 | 4.E-07 | 7.E-07 | 5.E-04 | 9.E-08 | 2.E-08 | 2.E-08 | 7.E-08 | 3.E-07 | 2.E-08 | 2.E-06 | 9.E-01 | 2.E-04 | 1.E-09 | 1.E-08 | 2.E-06 |
| 5xjh | 117 | LEU | 8.68% | 7.E-05 | 6.E-03 | 4.E-01 | 7.E-03 | 3.E-02 | 1.E-01 | 2.E-01 | 2.E-05 | 2.E-02 | 5.E-04 | 9.E-02 | 1.E-01 | 3.E-02 | 8.E-04 | 1.E-05 | 2.E-03 | 3.E-02 | 2.E-05 | 2.E-04 | 3.E-04 |
| 6ij6 | 104 | HIS | 9.20% | 7.E-09 | 1.E-06 | 2.E-05 | 1.E-06 | 1.E-07 | 1.E-05 | 7.E-06 | 3.E-10 | 9.E-02 | 4.E-08 | 5.E-08 | 7.E-07 | 4.E-06 | 6.E-02 | 6.E-10 | 4.E-08 | 2.E-08 | 5.E-04 | 9.E-01 | 3.E-08 |
| 5xjh | 213 | SER | 9.23% | 2.E-02 | 2.E-02 | 3.E-02 | 1.E-01 | 1.E-03 | 3.E-02 | 2.E-01 | 9.E-05 | 5.E-04 | 5.E-03 | 2.E-02 | 7.E-02 | 3.E-03 | 2.E-04 | 5.E-05 | 9.E-02 | 3.E-01 | 8.E-05 | 4.E-04 | 3.E-02 |
| 6ij6 | 240 | ALA | 9.40% | 9.E-02 | 2.E-05 | 1.E-03 | 9.E-05 | 8.E-01 | 3.E-04 | 1.E-04 | 1.E-05 | 1.E-05 | 3.E-04 | 7.E-05 | 4.E-05 | 2.E-04 | 1.E-06 | 3.E-05 | 7.E-03 | 6.E-02 | 8.E-07 | 9.E-07 | 2.E-02 |
| 6ij6 | 46 | SER | 9.48% | 8.E-01 | 7.E-03 | 3.E-03 | 3.E-02 | 1.E-03 | 7.E-03 | 6.E-02 | 4.E-03 | 1.E-04 | 5.E-05 | 5.E-04 | 3.E-02 | 5.E-04 | 3.E-05 | 9.E-05 | 9.E-02 | 3.E-03 | 1.E-05 | 3.E-05 | 3.E-04 |
| 6ij6 | 270 | THR | 10.14% | 2.E-05 | 2.E-03 | 2.E-03 | 6.E-04 | 3.E-04 | 5.E-03 | 4.E-03 | 3.E-07 | 7.E-05 | 2.E-01 | 1.E-03 | 1.E-02 | 8.E-04 | 4.E-06 | 2.E-06 | 1.E-04 | 1.E-01 | 1.E-06 | 3.E-06 | 6.E-01 |
| 6ij6 | 183 | ALA | 12.26% | 1.E-01 | 3.E-04 | 2.E-01 | 1.E-02 | 3.E-01 | 2.E-02 | 3.E-02 | 8.E-05 | 1.E-04 | 8.E-05 | 2.E-05 | 9.E-05 | 2.E-02 | 5.E-06 | 1.E-04 | 2.E-01 | 1.E-01 | 4.E-06 | 8.E-06 | 1.E-02 |
| 6ij6 | 110 | THR | 12.30% | 7.E-05 | 1.E-06 | 9.E-03 | 1.E-04 | 9.E-04 | 5.E-05 | 3.E-05 | 6.E-07 | 3.E-06 | 7.E-01 | 7.E-04 | 5.E-05 | 2.E-05 | 3.E-08 | 9.E-07 | 2.E-05 | 1.E-01 | 2.E-08 | 8.E-08 | 2.E-01 |
| 5xjh | 270 | THR | 12.72% | 5.E-03 | 1.E-04 | 3.E-03 | 6.E-04 | 7.E-03 | 4.E-03 | 3.E-04 | 3.E-05 | 3.E-05 | 3.E-02 | 1.E-03 | 5.E-04 | 6.E-04 | 8.E-06 | 6.E-03 | 5.E-03 | 1.E-01 | 8.E-07 | 6.E-06 | 8.E-01 |
| 6ij6 | 238 | SER | 13.39% | 8.E-01 | 1.E-07 | 2.E-05 | 8.E-05 | 8.E-05 | 1.E-07 | 1.E-07 | 9.E-02 | 8.E-08 | 4.E-10 | 6.E-09 | 2.E-07 | 3.E-08 | 7.E-09 | 1.E-07 | 1.E-01 | 8.E-06 | 1.E-09 | 4.E-09 | 4.E-09 |
| 5xjh | 124 | SER | 13.51% | 9.E-01 | 2.E-09 | 2.E-07 | 1.E-07 | 6.E-05 | 4.E-08 | 2.E-08 | 7.E-06 | 5.E-08 | 7.E-12 | 3.E-10 | 2.E-09 | 2.E-08 | 1.E-09 | 3.E-09 | 1.E-01 | 9.E-07 | 3.E-11 | 8.E-10 | 5.E-10 |
| 6ij6 | 95 | LYS | 13.62% | 5.E-07 | 9.E-04 | 1.E-03 | 1.E-04 | 3.E-05 | 2.E-01 | 6.E-01 | 2.E-09 | 9.E-04 | 3.E-05 | 6.E-02 | 1.E-01 | 3.E-03 | 7.E-06 | 1.E-08 | 5.E-06 | 9.E-06 | 3.E-07 | 3.E-06 | 4.E-06 |
| 5xjh | 274 | GLU | 14.23% | 2.E-06 | 1.E-03 | 3.E-03 | 3.E-04 | 9.E-04 | 5.E-02 | 1.E-01 | 8.E-08 | 1.E-02 | 3.E-03 | 7.E-02 | 3.E-02 | 6.E-01 | 8.E-02 | 6.E-07 | 8.E-06 | 7.E-04 | 4.E-05 | 6.E-03 | 8.E-03 |
| 6ij6 | 119 | GLN | 14.67% | 8.E-07 | 2.E-02 | 7.E-05 | 3.E-05 | 1.E-05 | 1.E-01 | 2.E-03 | 5.E-06 | 5.E-03 | 2.E-05 | 1.E-02 | 1.E-01 | 2.E-01 | 3.E-01 | 2.E-07 | 9.E-07 | 2.E-06 | 1.E-02 | 2.E-01 | 1.E-06 |
| 5xjh | 51 | THR | 15.48% | 3.E-02 | 8.E-03 | 4.E-02 | 5.E-02 | 3.E-03 | 6.E-02 | 5.E-01 | 3.E-04 | 3.E-04 | 7.E-05 | 5.E-03 | 9.E-02 | 3.E-03 | 4.E-05 | 1.E-05 | 2.E-02 | 2.E-01 | 8.E-06 | 3.E-05 | 2.E-03 |
| 5xjh | 41 | ALA | 16.64% | 2.E-01 | 1.E-02 | 1.E-02 | 4.E-02 | 9.E-04 | 3.E-02 | 1.E-01 | 5.E-01 | 5.E-04 | 6.E-05 | 4.E-03 | 4.E-02 | 2.E-03 | 3.E-04 | 5.E-04 | 5.E-02 | 6.E-03 | 7.E-05 | 2.E-04 | 2.E-04 |
| 5xjh | 92 | SER | 17.08% | 8.E-01 | 2.E-03 | 9.E-04 | 1.E-02 | 2.E-04 | 2.E-03 | 2.E-02 | 4.E-05 | 7.E-06 | 1.E-05 | 3.E-05 | 2.E-03 | 3.E-05 | 2.E-06 | 5.E-05 | 2.E-01 | 1.E-03 | 1.E-06 | 4.E-06 | 1.E-04 |
| 5xjh | 63 | TYR | 17.23% | 4.E-09 | 7.E-06 | 5.E-07 | 7.E-09 | 2.E-07 | 2.E-06 | 8.E-08 | 1.E-09 | 5.E-03 | 3.E-09 | 3.E-07 | 9.E-07 | 7.E-05 | 8.E-01 | 2.E-10 | 1.E-08 | 1.E-08 | 2.E-04 | 2.E-01 | 8.E-09 |
| 5xjh | 133 | GLN | 17.50% | 4.E-06 | 3.E-05 | 4.E-02 | 2.E-02 | 1.E-04 | 2.E-01 | 7.E-01 | 1.E-07 | 8.E-04 | 2.E-03 | 6.E-03 | 1.E-03 | 1.E-03 | 2.E-05 | 8.E-07 | 1.E-05 | 1.E-02 | 2.E-06 | 1.E-05 | 8.E-03 |
| 6ij6 | 236 | SER | 18.79% | 1.E-01 | 3.E-04 | 7.E-02 | 6.E-01 | 1.E-03 | 6.E-04 | 3.E-04 | 4.E-04 | 1.E-05 | 7.E-06 | 1.E-05 | 4.E-04 | 3.E-05 | 7.E-07 | 1.E-05 | 2.E-01 | 5.E-04 | 1.E-06 | 9.E-07 | 3.E-05 |
| 5xjh | 292 | GLU | 19.20% | 4.E-03 | 3.E-02 | 1.E-01 | 7.E-02 | 4.E-03 | 8.E-02 | 2.E-01 | 9.E-04 | 6.E-02 | 2.E-02 | 1.E-01 | 2.E-01 | 2.E-02 | 2.E-02 | 3.E-04 | 3.E-02 | 3.E-02 | 1.E-03 | 5.E-03 | 1.E-02 |

| | | | | | | | | | | | | | | | | | | | | | | | |
|---|---|---|---|---|---|---|---|---|---|---|---|---|---|---|---|---|---|---|---|---|---|---|---|
| 5xjh | 208 | ILE | 19.53% | 6.E-04 | 9.E-03 | 3.E-03 | 2.E-03 | 2.E-03 | 1.E-02 | 2.E-02 | 4.E-05 | 3.E-04 | 2.E-01 | 1.E-03 | 1.E-02 | 5.E-03 | 2.E-04 | 5.E-05 | 2.E-03 | 2.E-01 | 1.E-04 | 1.E-04 | 6.E-01 |
| 6ij6 | 269 | SER | 20.31% | 5.E-01 | 3.E-04 | 6.E-02 | 1.E-01 | 8.E-03 | 1.E-02 | 7.E-03 | 3.E-05 | 2.E-03 | 1.E-03 | 2.E-04 | 2.E-03 | 2.E-04 | 2.E-05 | 7.E-06 | 2.E-01 | 4.E-02 | 4.E-06 | 3.E-05 | 8.E-03 |
| 6ij6 | 190 | ASN | 21.04% | 5.E-07 | 8.E-06 | 2.E-01 | 8.E-01 | 3.E-05 | 4.E-05 | 1.E-04 | 3.E-08 | 4.E-07 | 3.E-07 | 5.E-07 | 2.E-05 | 1.E-06 | 2.E-08 | 2.E-08 | 4.E-05 | 1.E-05 | 1.E-08 | 1.E-08 | 2.E-06 |
| 6ij6 | 41 | ALA | 22.07% | 2.E-01 | 4.E-02 | 3.E-02 | 6.E-02 | 6.E-04 | 9.E-02 | 4.E-01 | 1.E-04 | 9.E-03 | 6.E-04 | 6.E-03 | 1.E-01 | 2.E-03 | 1.E-03 | 6.E-05 | 3.E-02 | 3.E-02 | 5.E-04 | 1.E-03 | 2.E-03 |
| 5xjh | 154 | MET | 22.55% | 5.E-07 | 5.E-03 | 4.E-04 | 1.E-05 | 1.E-05 | 7.E-01 | 5.E-02 | 5.E-08 | 1.E-04 | 5.E-06 | 1.E-03 | 3.E-02 | 2.E-01 | 4.E-04 | 3.E-08 | 2.E-07 | 7.E-07 | 2.E-04 | 2.E-04 | 2.E-07 |
| 5xjh | 207 | SER | 22.65% | 8.E-02 | 1.E-02 | 2.E-01 | 4.E-01 | 8.E-03 | 8.E-03 | 4.E-02 | 5.E-04 | 5.E-03 | 2.E-04 | 5.E-04 | 6.E-03 | 2.E-03 | 6.E-04 | 7.E-05 | 2.E-01 | 1.E-02 | 8.E-05 | 7.E-04 | 1.E-03 |
| 5xjh | 73 | ASN | 22.79% | 5.E-06 | 4.E-06 | 2.E-01 | 8.E-01 | 2.E-04 | 2.E-04 | 2.E-03 | 2.E-07 | 3.E-05 | 3.E-06 | 3.E-06 | 4.E-05 | 3.E-06 | 2.E-07 | 4.E-07 | 4.E-04 | 2.E-05 | 3.E-08 | 2.E-07 | 1.E-05 |
| 6ij6 | 207 | SER | 23.23% | 2.E-01 | 1.E-02 | 1.E-01 | 4.E-01 | 4.E-03 | 5.E-03 | 1.E-02 | 2.E-03 | 7.E-03 | 4.E-04 | 5.E-04 | 8.E-03 | 8.E-04 | 2.E-03 | 9.E-05 | 2.E-01 | 7.E-03 | 9.E-04 | 2.E-03 | 2.E-03 |
| 6ij6 | 261 | PHE | 23.26% | 1.E-08 | 4.E-06 | 1.E-03 | 4.E-07 | 6.E-07 | 9.E-05 | 4.E-05 | 3.E-09 | 4.E-01 | 8.E-08 | 1.E-07 | 8.E-06 | 1.E-04 | 2.E-01 | 2.E-09 | 2.E-08 | 2.E-08 | 3.E-01 | 1.E-01 | 3.E-09 |
| 5xjh | 190 | ASN | 23.32% | 2.E-05 | 4.E-03 | 2.E-01 | 8.E-01 | 2.E-04 | 1.E-03 | 2.E-03 | 4.E-06 | 6.E-04 | 7.E-06 | 1.E-04 | 2.E-03 | 2.E-04 | 9.E-05 | 1.E-06 | 6.E-04 | 1.E-04 | 1.E-04 | 9.E-05 | 4.E-05 |
| 6ij6 | 165 | GLY | 25.56% | 7.E-01 | 1.E-07 | 2.E-07 | 3.E-07 | 1.E-05 | 7.E-08 | 5.E-08 | 3.E-01 | 2.E-08 | 3.E-09 | 2.E-08 | 8.E-08 | 3.E-08 | 1.E-08 | 8.E-06 | 4.E-03 | 1.E-06 | 1.E-09 | 3.E-09 | 4.E-08 |
| 6ij6 | 125 | SER | 26.36% | 9.E-02 | 3.E-03 | 1.E-03 | 4.E-03 | 6.E-03 | 1.E-02 | 2.E-02 | 1.E-04 | 3.E-04 | 2.E-03 | 2.E-03 | 1.E-02 | 1.E-03 | 2.E-04 | 6.E-05 | 3.E-01 | 4.E-01 | 5.E-05 | 2.E-04 | 2.E-01 |
| 6ij6 | 175 | SER | 26.87% | 2.E-01 | 4.E-03 | 4.E-02 | 4.E-01 | 1.E-03 | 3.E-03 | 4.E-02 | 4.E-04 | 4.E-04 | 1.E-05 | 8.E-04 | 9.E-03 | 5.E-04 | 8.E-05 | 5.E-04 | 3.E-01 | 1.E-02 | 1.E-05 | 8.E-05 | 8.E-05 |
| 5xjh | 37 | ASN | 27.55% | 7.E-07 | 2.E-04 | 3.E-01 | 7.E-01 | 9.E-05 | 7.E-05 | 1.E-03 | 7.E-08 | 1.E-03 | 7.E-04 | 2.E-05 | 1.E-03 | 4.E-05 | 2.E-05 | 4.E-07 | 4.E-05 | 7.E-04 | 7.E-07 | 1.E-05 | 9.E-04 |
| 5xjh | 287 | ALA | 27.60% | 3.E-01 | 4.E-03 | 9.E-03 | 2.E-01 | 7.E-03 | 4.E-03 | 2.E-02 | 2.E-03 | 4.E-03 | 2.E-05 | 2.E-04 | 2.E-02 | 1.E-03 | 3.E-04 | 3.E-05 | 5.E-01 | 6.E-03 | 1.E-04 | 3.E-04 | 8.E-04 |
| 6ij6 | 63 | TYR | 27.81% | 8.E-09 | 5.E-06 | 3.E-06 | 2.E-08 | 6.E-07 | 1.E-05 | 2.E-07 | 3.E-09 | 2.E-02 | 8.E-09 | 1.E-06 | 1.E-06 | 2.E-04 | 7.E-01 | 6.E-10 | 3.E-08 | 5.E-08 | 3.E-04 | 3.E-01 | 2.E-08 |
| 6ij6 | 231 | GLU | 28.31% | 3.E-06 | 3.E-04 | 3.E-03 | 3.E-05 | 5.E-03 | 7.E-01 | 3.E-01 | 4.E-08 | 2.E-06 | 2.E-06 | 1.E-03 | 5.E-03 | 8.E-03 | 1.E-08 | 1.E-08 | 9.E-05 | 1.E-06 | 6.E-08 | 9.E-09 | 5.E-07 |
| 6ij6 | 133 | GLN | 28.50% | 3.E-06 | 2.E-04 | 4.E-02 | 1.E-01 | 9.E-05 | 3.E-01 | 5.E-01 | 3.E-07 | 5.E-03 | 6.E-04 | 5.E-03 | 4.E-03 | 2.E-03 | 7.E-05 | 1.E-06 | 1.E-05 | 1.E-03 | 8.E-06 | 7.E-05 | 6.E-03 |
| 5xjh | 159 | TRP | 28.75% | 1.E-10 | 3.E-05 | 3.E-07 | 1.E-07 | 9.E-09 | 8.E-06 | 4.E-07 | 5.E-11 | 3.E-02 | 3.E-10 | 9.E-07 | 3.E-05 | 3.E-05 | 3.E-01 | 5.E-11 | 6.E-10 | 3.E-10 | 3.E-01 | 3.E-01 | 4.E-10 |
| 6ij6 | 132 | ARG | 29.00% | 3.E-05 | 3.E-01 | 4.E-03 | 1.E-03 | 8.E-05 | 4.E-01 | 3.E-02 | 7.E-07 | 6.E-02 | 7.E-05 | 5.E-04 | 8.E-02 | 1.E-02 | 3.E-02 | 1.E-06 | 4.E-05 | 4.E-04 | 6.E-02 | 2.E-02 | 2.E-04 |
| 5xjh | 175 | SER | 29.89% | 5.E-01 | 8.E-03 | 1.E-02 | 1.E-01 | 2.E-03 | 1.E-02 | 4.E-02 | 1.E-04 | 3.E-04 | 2.E-05 | 2.E-03 | 3.E-02 | 8.E-04 | 8.E-05 | 1.E-05 | 3.E-01 | 1.E-02 | 1.E-05 | 7.E-05 | 3.E-04 |

**Supplementary Information Fig. 2 | Disfavored PETase residues flagged by MutCompute from the wild-type and ThermoPETase crystal structures.** MutCompute outputs a probability distribution that describes the likelihood of each of the 20 canonical amino acids to be the wild-type amino acid for the surrounding chemical environment. A disfavored residue is defined as a residue where the amino acid with the highest predicted probability is not the wild-type amino acid. Here, a 30% wild-type probability cutoff was used to down select disfavored residues.

| Ranking | position | wtAA | prAA | wt_prob | pred_prob | avg_log_ratio |
|---|---|---|---|---|---|---|
| 1 | 121 | SER | GLU | 0.11% | 61.20% | 6.86 |
| 2 | 262 | MET | LEU | 4.12% | 66.51% | 4.77 |
| 3 | 233 | ASN | LYS | 4.86% | 55.93% | 3.06 |
| 4 | 140 | THR | ASP | 14.23% | 75.20% | 2.69 |
| 5 | 58 | SER | GLU | 5.22% | 45.81% | 2.49 |
| 6 | 169 | SER | ALA | 10.29% | 89.60% | 2.46 |
| 7 | 119 | GLN | LEU | 6.06% | 54.65% | 2.40 |
| 8 | 225 | ASN | CYS | 7.94% | 78.08% | 2.31 |
| 9 | 270 | THR | VAL | 8.65% | 72.13% | 2.26 |
| 10 | 114 | ASN | THR | 9.53% | 76.00% | 2.14 |
| 11 | 91 | GLN | ILE | 10.03% | 53.89% | 2.14 |
| 12 | 207 | SER | ASP | 15.83% | 37.06% | 1.78 |
| 13 | 212 | ASN | ALA | 6.17% | 33.75% | 1.73 |
| 14 | 59 | ARG | ASN | 6.26% | 34.64% | 1.69 |
| 15 | 136 | SER | LEU | 5.99% | 28.02% | 1.66 |
| 16 | 279 | THR | GLU | 6.85% | 26.04% | 1.66 |
| 17 | 168 | ILE | LEU | 28.37% | 71.44% | 1.65 |
| 18 | 263 | ASP | ASN | 19.55% | 32.91% | 1.65 |
| 19 | 154 | MET | GLN | 10.19% | 46.02% | 1.64 |
| 20 | 190 | ASN | ASP | 21.64% | 77.84% | 1.62 |
| 21 | 201 | PHE | LEU | 24.70% | 67.93% | 1.61 |
| 22 | 124 | SER | ALA | 25.46% | 74.53% | 1.32 |
| 23 | 208 | ILE | VAL | 18.96% | 49.48% | 1.23 |
| 24 | 117 | LEU | LYS | 9.84% | 25.37% | 1.21 |
| 25 | 95 | LYS | GLU | 20.94% | 44.44% | 1.11 |
| 26 | 73 | ASN | ASP | 28.55% | 70.65% | 1.06 |
| 27 | 53 | ARG | LYS | 29.40% | 66.87% | 1.00 |
| 28 | 274 | GLU | MET | 22.29% | 46.47% | 0.99 |
| 29 | 67 | THR | VAL | 29.74% | 50.17% | 0.97 |
| 30 | 125 | SER | ALA | 18.70% | 38.47% | 0.59 |
| 31 | 213 | SER | THR | 27.53% | 40.39% | 0.35 |
| 32 | 146 | TYR | HIS | 27.67% | 44.27% | 0.34 |
| 33 | 88 | THR | THR | 22.54% | 22.54% | 0.00 |
| 34 | 172 | ASN | ASN | 25.92% | 25.92% | 0.00 |
| 35 | 187 | SER | SER | 27.74% | 27.74% | 0.00 |
| 36 | 292 | GLU | GLU | 29.43% | 29.43% | 0.00 |
| 37 | 152 | ALA | GLY | 22.79% | 35.05% | -0.10 |

**Supplementary Information Fig. 3a | Predictions (based on wild-type PETase) ranked by fold change in the probabilities between the predicted and the wild-type amino acid.** Fold change predictions are provided as a means of down-selecting potential mutations.

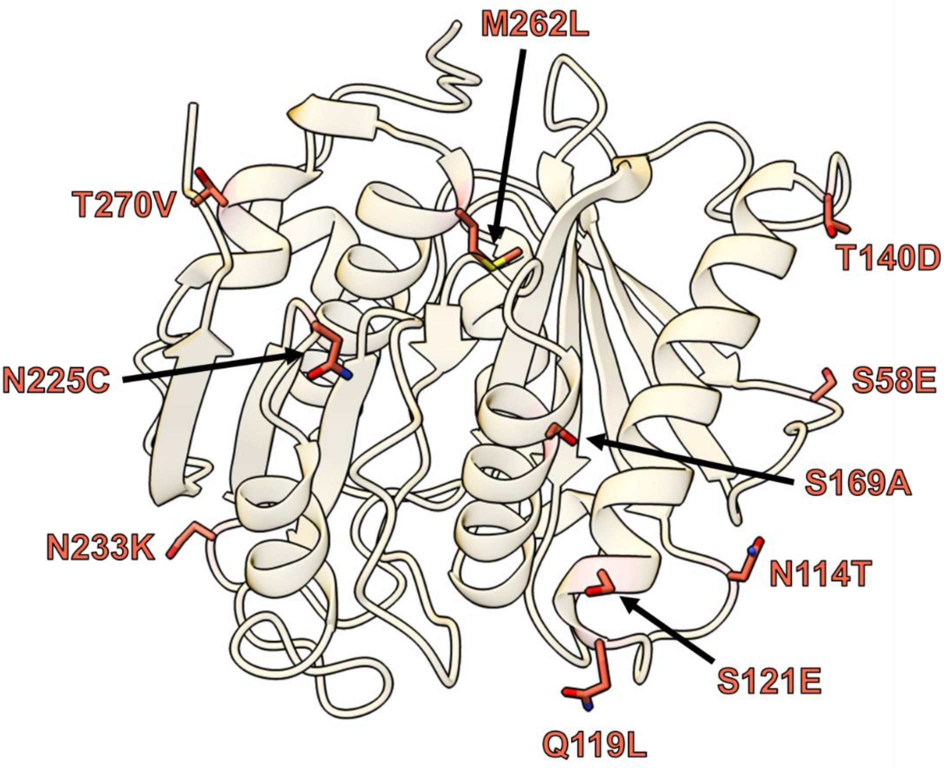


**Supplementary Information Fig. 3b | TOP 10 ranked predictions (based on wild-type PETase).** The top 10 mutations predicted for the wild-type PETase scaffold are presented.

| Ranking | position | wtAA | prAA | wt_prob | pred_prob | avg_log_ratio |
|---|---|---|---|---|---|---|
| 1 | 91 | GLN | ILE | 0.09% | 98.29% | 7.42 |
| 2 | 233 | ASN | LYS | 0.09% | 97.29% | 7.39 |
| 3 | 225 | ASN | CYS | 0.32% | 98.01% | 5.86 |
| 4 | 61 | SER | THR | 0.54% | 31.85% | 3.78 |
| 5 | 208 | ILE | VAL | 3.02% | 88.68% | 3.57 |
| 6 | 58 | SER | ALA | 2.30% | 78.28% | 3.54 |
| 7 | 34 | ARG | LEU | 2.99% | 78.89% | 3.41 |
| 8 | 240 | ALA | CYS | 9.40% | 81.59% | 3.24 |
| 9 | 186 | HIS | ASP | 4.75% | 77.21% | 3.20 |
| 10 | 224 | ARG | GLN | 3.52% | 39.61% | 3.14 |
| 11 | 53 | ARG | LYS | 2.82% | 56.07% | 2.99 |
| 12 | 270 | THR | VAL | 10.14% | 63.14% | 2.91 |
| 13 | 136 | SER | THR | 4.15% | 66.37% | 2.82 |
| 14 | 37 | ASN | ASP | 6.92% | 92.96% | 2.65 |
| 15 | 104 | HIS | TYR | 9.20% | 85.21% | 2.33 |
| 16 | 46 | SER | ALA | 9.48% | 75.76% | 2.31 |
| 17 | 95 | LYS | GLU | 13.62% | 57.80% | 2.22 |
| 18 | 59 | ARG | THR | 3.25% | 31.60% | 2.18 |
| 19 | 87 | TYR | ARG | 8.43% | 50.29% | 2.14 |
| 20 | 238 | SER | ALA | 13.39% | 78.03% | 2.09 |
| 21 | 236 | SER | ASP | 18.79% | 61.12% | 1.82 |
| 22 | 110 | THR | ILE | 12.30% | 69.39% | 1.75 |
| 23 | 33 | MET | GLN | 7.37% | 36.75% | 1.69 |
| 24 | 165 | GLY | ALA | 25.56% | 74.08% | 1.55 |
| 25 | 279 | THR | GLU | 7.58% | 31.98% | 1.44 |
| 26 | 119 | GLN | PHE | 14.67% | 29.22% | 1.35 |
| 27 | 190 | ASN | ASP | 21.04% | 78.93% | 1.33 |
| 28 | 63 | TYR | PHE | 27.81% | 69.99% | 1.32 |
| 29 | 212 | ASN | ALA | 8.26% | 23.24% | 1.10 |
| 30 | 269 | SER | ALA | 20.31% | 54.77% | 0.98 |
| 31 | 231 | GLU | GLN | 28.31% | 69.43% | 0.93 |
| 32 | 133 | GLN | GLU | 28.50% | 52.67% | 0.80 |
| 33 | 183 | ALA | CYS | 12.26% | 30.00% | 0.74 |
| 34 | 175 | SER | ASP | 26.87% | 42.64% | 0.71 |
| 35 | 41 | ALA | GLU | 22.07% | 38.13% | 0.55 |
| 36 | 261 | PHE | HIS | 23.26% | 37.42% | 0.52 |
| 37 | 125 | SER | THR | 26.36% | 39.71% | 0.47 |
| 38 | 207 | SER | ASP | 23.23% | 44.12% | 0.46 |
| 39 | 132 | ARG | GLN | 29.00% | 41.70% | 0.37 |
| 40 | 277 | ASN | ASN | 23.99% | 23.99% | 0.00 |

**Supplementary Information Fig. 3c | Predictions (based on ThermoPETase) ranked by fold change in the probabilities between the predicted and the wild-type amino acid.** Fold change predictions are provided as a means of down-selecting potential mutations.

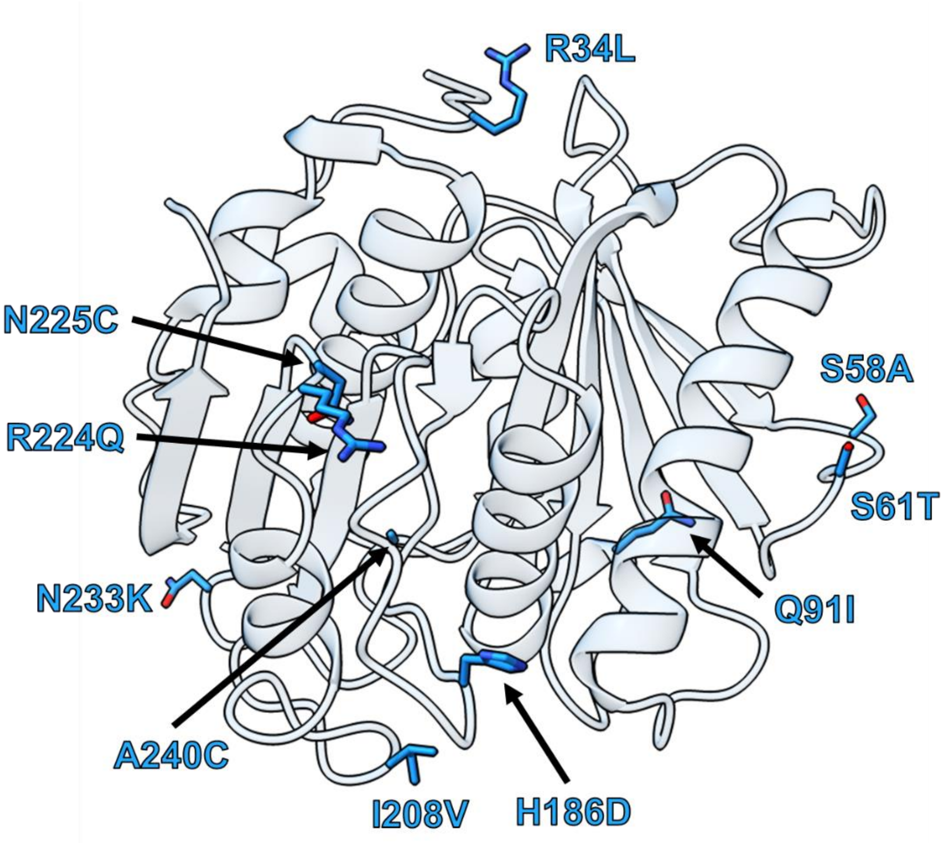


**Supplementary Information Fig. 3d | TOP 10 ranked predictions (based on ThermoPETase).** The top 10 mutations predicted for the ThermoPETase scaffold are presented.

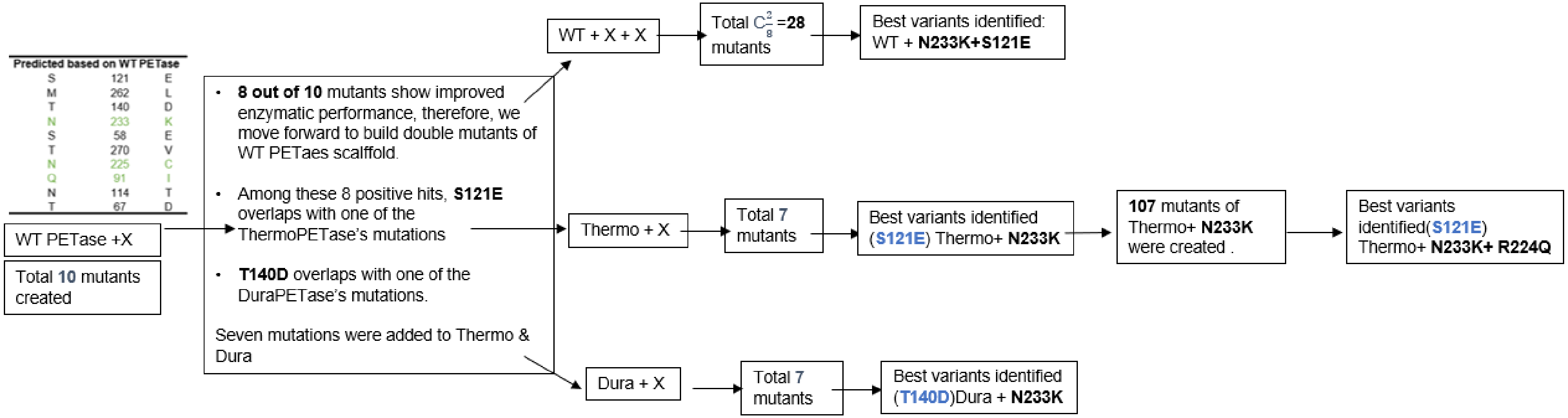


**Supplementary Information Fig. 4 | Selecting mutations based on experimental catalytic activity measurements.** A scheme for selecting mutations based on experimental evidence is provided.

Initially, we chose the top ten ranked predictions based on crystal structure PDB: 5XJH (Supplementary Information Fig. 3a) and introduced them respectively into the wild-type PETase scaffold to generate ten single mutants. Experimental characterization of these variants showed that eight out of these ten predicted mutations confer improved thermostability and activity to the wild-type PETase scaffold. Notably, of such eight beneficial mutations, S121E and T140D are each overlapped with one of the mutations of ThermoPETase and DuraPETase.

Subsequently, we paired two of the eight beneficial mutations to create all 28 possible double mutants of wild-type PETase. Meanwhile, the unique beneficial mutations were respectively introduced into ThermoPETase and DuraPETase scaffolds to generate 14 variants that contain two predictions from Mutcompute. Among these 42 variants, the best variants identified for each scaffold all contain the predicted mutation-N233K. As the variant exhibiting the highest enzymatic activity, ThermoPETase$^{N233K}$ was chosen as the template for further mutagenesis.

Finally, a total of 107 variants of ThermoPETase$^{N233K}$ were created by incorporating single or multiple mutations from the 14 top ranked predictions based on both crystal structures PDB: 5XJH and 6IJ6 as well as lower ranked predictions selected by a rational design strategy. After comparative analysis of the enzymatic performance of these 107 variants, ThermoPETase$^{N233K/R224Q}$ was identified as the best variant as it showed improved activity versus ThermoPETase$^{N233K}$. Given the synergetic interactions among the four predicted mutations that resulted in the best WT PETase, ThermoPETase and DuraPETase variants, S121E, N233K, R224Q and T140D were further selected for combinatorial assembly and follow-up analysis.

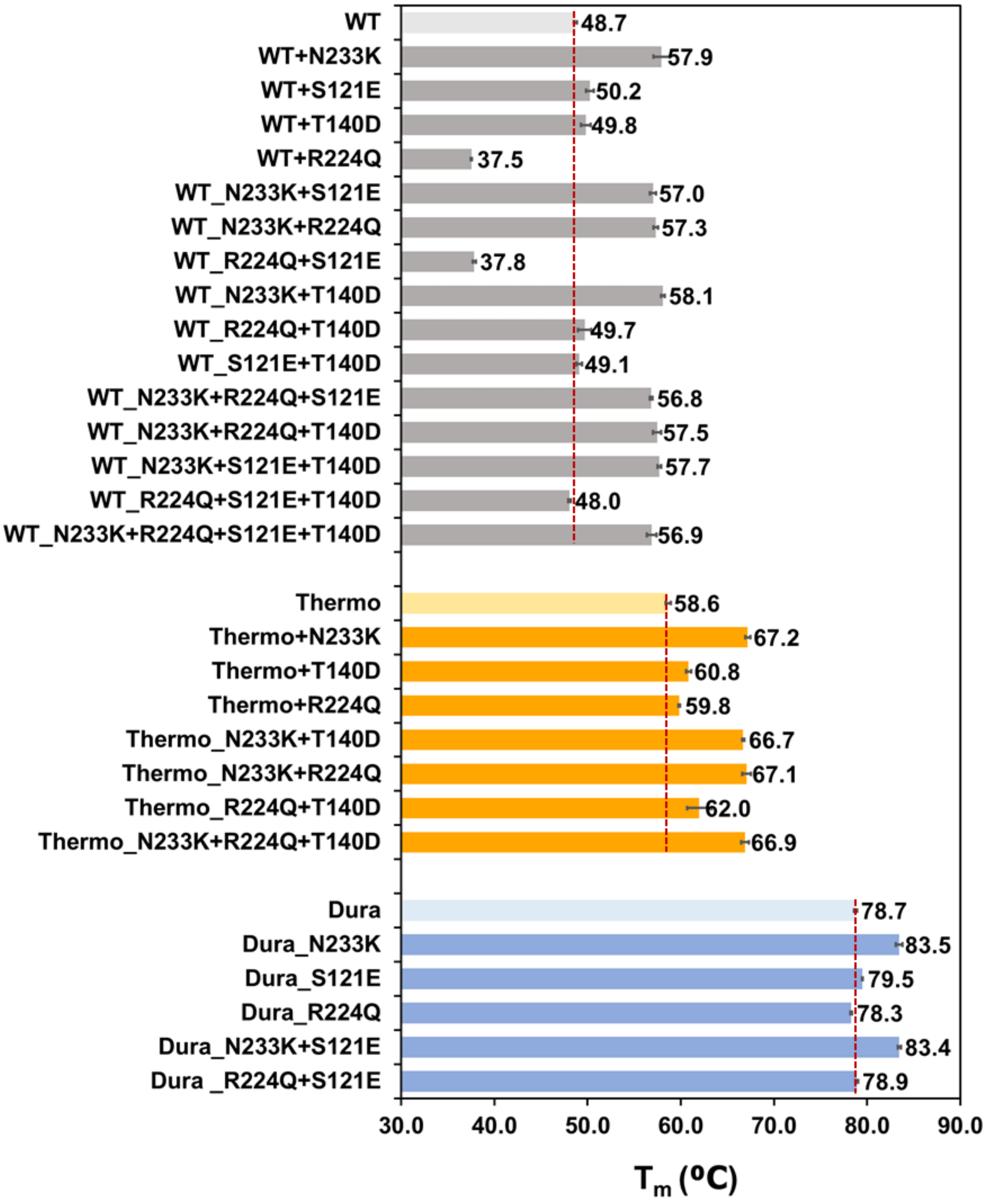


**Supplementary Information Fig. 5 | Thermostability of the PETase variants incorporating the mutations predicted by Mutcompute and their respective scaffolds—wild-type PETase (WT), ThermoPETase (Thermo), DuraPETase (Dura).** The melting temperature of each enzyme was determined by DSC. All measurement were conducted in triplicate (n=3).

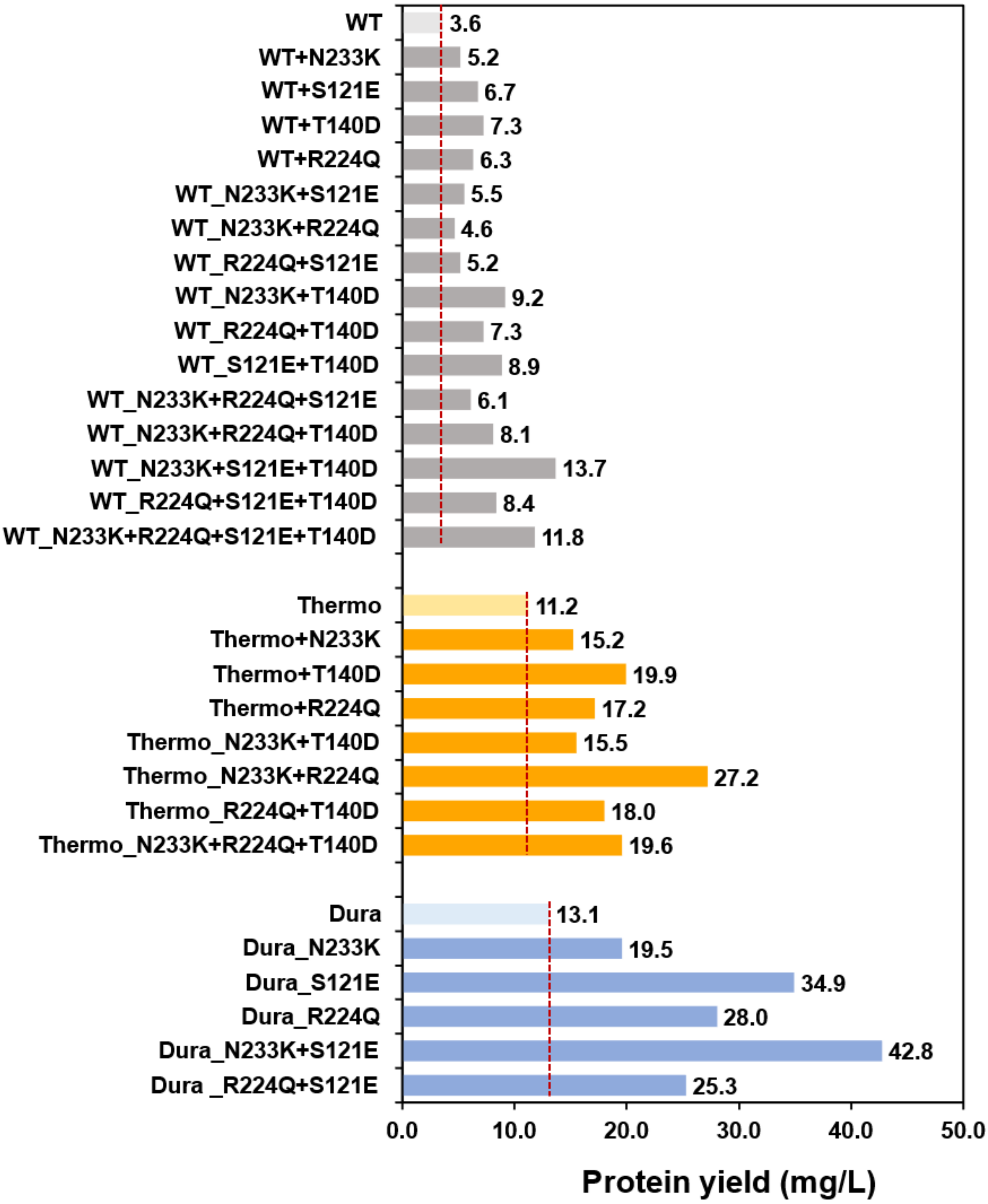


**Supplementary Information Fig. 6 | Protein yield of the PETase variants incorporating the mutations predicted by Mutcompute and their respective scaffolds—wild-type PETase (WT), ThermoPETase (Thermo), DuraPETase (Dura).** Protein yields from *P. putida* purification experiments indicate improved yields from mutant enzymes.

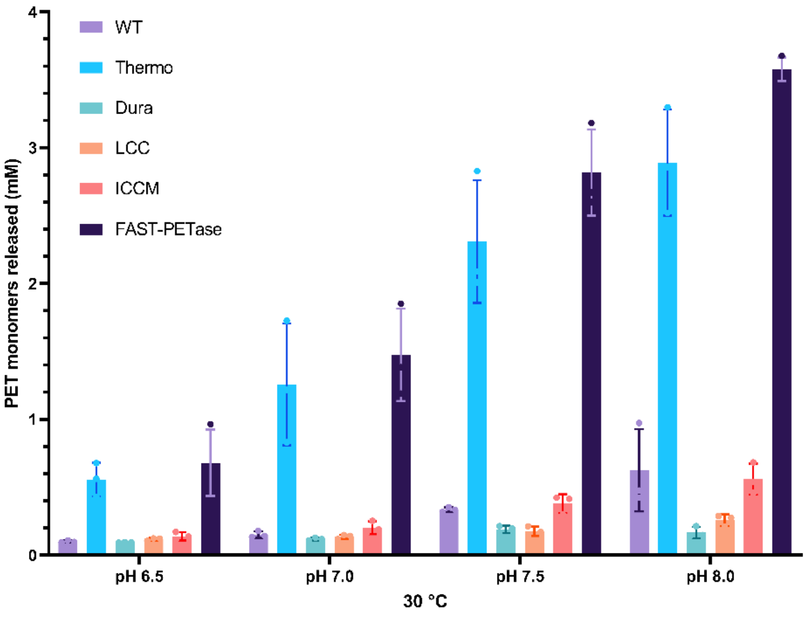


**a**

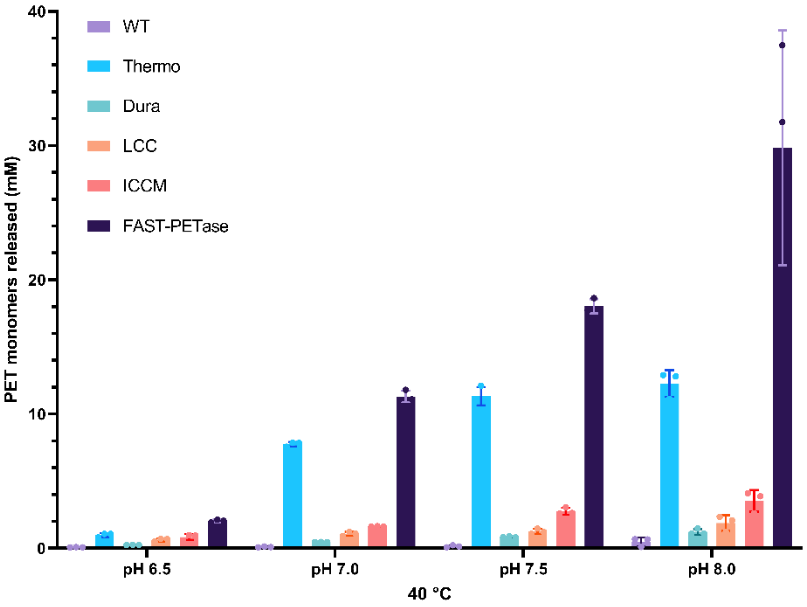


**b**

**Supplementary Information Fig. 7 | The PET-hydrolytic activity of FAST-PETase outperformed various PHEs at mild temperatures and modest pH.** Comparison of PET-hydrolytic activity of FAST-PETase, wild-type PETase (WT), ThermoPETase (Thermo), DuraPETase (Dura), LCC and ICCM across a range of pH (6.5 – 8.0) at reaction temperatures of 30 ºC (**a.**) and 40 ºC (**b.**). PET-hydrolytic activity was evaluated by measuring the amount of PET monomers (the sum of TPA and MHET) released from hydrolyzing gf-PET film by the tested enzymes after 96 hrs of reaction time. All measurement were conducted in triplicate (n=3).

| Sample number | Postconsumer Plastic products | Initial mass (mg) | Crystallinity % | Time for complete degradation (days) | Category | Mn kg/mol | Mw kg/mol | Đ |
|---|---|---|---|---|---|---|---|---|
| #1 | 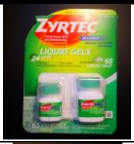 | 9.49 ± 0.27 | 1.18% ± 0.02％ | 2.5 | Medication packaging | 30.2 | 55.4 | 1.83 |
| #2 | 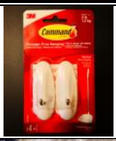 | 6.36 ± 0.07 | 1.21％ ± 0.09％ | 2.5 | Household goods packaging | 31.6 | 56.2 | 1.78 |
| #3 | 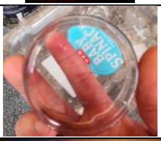 | 16.27 ± 0.52 | 1.23％ ± 0.20％ | 4.5 | Beverage packaging | 30.7 | 53.9 | 1.76 |
| #4 | 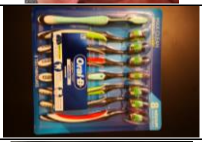 | 12.34 ± 0.11 | 1.30％ ± 0.14％ | 4 | Household goods packaging | 33.1 | 61.2 | 1.85 |
| #5 | 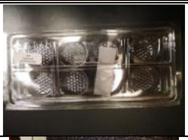 | 8.28 ± 0.48 | 1.40％ ± 0.14％ | 2 | Food packaging | 29.2 | 50.9 | 1.74 |
| #6 | 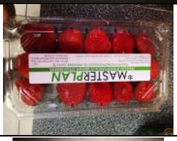 | 15.75 ± 0.11 | 1.42％ ± 0.29％ | 4.5 | Food packaging | 29.0 | 50.9 | 1.76 |
| #7 | 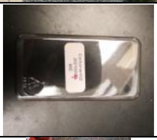 | 4.86 ± 0.32 | 1.44％ ± 0.25％ | 2 | Household goods packaging | 29.8 | 57.9 | 1.94 |
| #8 | 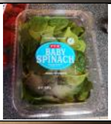 | 10.57 ± 0.02 | 1.50％ ± 0.21％ | 3.5 | Food packaging | 33.2 | 59.9 | 1.80 |
| #9 | 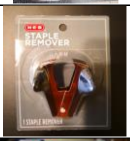 | 6.41 ± 0.23 | 1.54％ ± 0.13％ | 2.5 | Office supplies packaging | 32.1 | 56.8 | 1.77 |
| #10 | 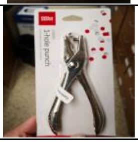 | 8.76 ± 0.24 | 1.55％ ± 0.22％ | 2 | Office supplies packaging | 35.4 | 66.6 | 1.88 |

**Supplementary Information Fig. 8 | Mass, crystallinity %, molecular weights (Mn, Mw), polydispersity indices (Đ) and time for complete degradation of various pc-PET films by FAST-PETase.** The circular pc-PET films (6 mm in diameter) were hole-punched from 51 different post-consumer plastic products used in the packaging of food, beverages, medications, office supplies, household goods and cosmetics available at local grocery store chains (Walmart, Costco, and HEB). The pc-PET films were hydrolysed by serial treatment with FAST-PETase at 50 ºC until the films were completed degraded. The enzyme solution (200 nM of FAST-PETase in 100mM $KH_2PO4$-NaOH (pH 8.0) buffer) was replenished every 24 hours. The crystallinity % of the intact pc-PET films was determined by DSC. The initial mass of the films was determined gravimetrically by a digital scale. Both DSC and gravimetric measurements were conducted in triplicate. Means ± s.d. (n=3) are shown.

| Sample number | Postconsumer Plastic products | Initial mass (mg) | Crystallinity % | Time for complete degradation (days) | Category | Mn kg/mol | Mw kg/mol | Đ |
|---|---|---|---|---|---|---|---|---|
| #11 | 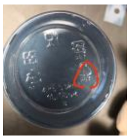 | 9.46 ± 0.2 | 1.65% ± 0.08% | 2.5 | Food packaging | 33.3 | 60.6 | 1.82 |
| #12 | 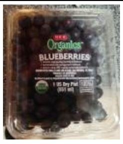 | 10.20 ± 0.5 | 1.65% ± 0.21% | 2.5 | Cosmetics packaging | 27.2 | 50.3 | 1.85 |
| #13 | 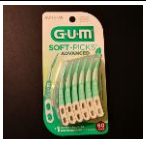 | 11.54 ± 0.27 | 1.68% ± 0.30% | 4 | Food packaging | 32.6 | 59.8 | 1.83 |
| #14 | 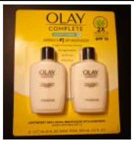 | 15.44 ± 0.25 | 1.68% ± 0.06% | 5 | Food packaging | 27.1 | 48.2 | 1.78 |
| #15 | 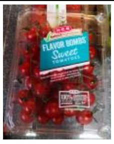 | 13.06 ± 0.72 | 1.73% ± 0.27% | 2.5 | Medication packaging | 28.2 | 52.3 | 1.85 |
| #16 | 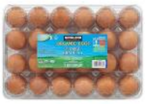 | 11.84 ± 1.06 | 2.00% ± 0.16% | 5 | Medication packaging | 23.3 | 45.3 | 1.94 |
| #17 | 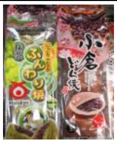 | 4.37 ± 0.66 | 2.01% ± 0.03% | 1 | Food packaging | 30.9 | 54.7 | 1.77 |
| #18 | 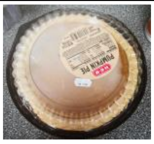 | 17.13 ± 0.17 | 2.14% ± 0.21% | 4.5 | Office supplies packaging | 36.4 | 62.4 | 1.71 |
| #19 | 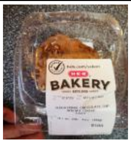 | 10.83 ± 0.69 | 2.19% ± 0.22% | 1.5 | Household goods packaging | 36.1 | 62.5 | 1.73 |
| #20 | 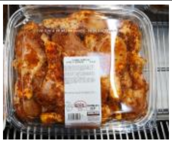 | 11.25 ± 0.25 | 2.19% ± 0.07% | 3.5 | Food packaging | 26.0 | 52.4 | 2.02 |
| #21 | 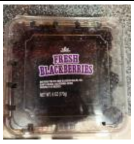 | 14.34 ± 0.1 | 2.20% ± 0.17% | 3.5 | Household goods packaging | 24.0 | 44.1 | 1.84 |

Supplementary Information Fig. 8 continued

| Sample number | Postconsumer Plastic products | Initial mass (mg) | Crystallinity % | Time for complete degradation (days) | Category | Mn kg/mol | Mw kg/mol | Đ |
|---|---|---|---|---|---|---|---|---|
| **#22** | 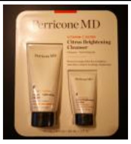  | 17.06 ± 0.74 | 2.28% ± 0.24% | 7 | Cosmetics packaging | 36.2 | 66.3 | 1.83 |
| **#23** | 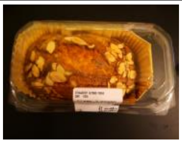 | 11.27 ± 0.71 | 2.29% ± 0.13% | 4 | Food packaging | 24.8 | 61.5 | 2.48 |
| **#24** | 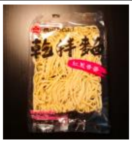 | 7.07 ± 0.12 | 2.45% ± 0.17% | 2.5 | Food packaging | 28.1 | 50.9 | 1.81 |
| **#25** | 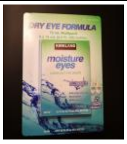 | 11.06 ± 0.08 | 2.49% ± 0.26% | 5 | Medication packaging | 33.4 | 61.1 | 1.83 |
| **#26** | 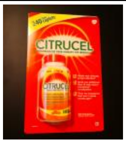  | 9.34 ± 0.28 | 2.53% ± 0.22% | 4 | Medication packaging | 32.7 | 57.8 | 1.77 |
| **#27** | 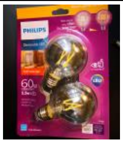 | 10.52 ± 0.07 | 2.53% ± 1.64% | 5 | Household goods packaging | 32.7 | 59.9 | 1.83 |
| **#28** | 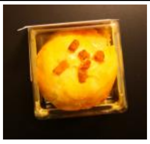 | 8.97 ± 0.37 | 2.56% ± 0.16% | 4 | Food packaging | 28.7 | 53.7 | 1.87 |
| **#29** | 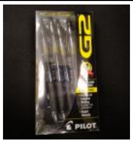  | 13.04 ± 0.03 | 2.56% ± 0.24% | 6 | Office supplies packaging | 33.7 | 62.3 | 1.85 |
| **#30** | 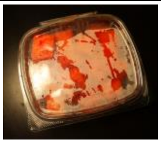 | 8.16 ± 2.23 | 2.61% ± 1.05% | 3.5 | Food packaging | 38.0 | 70.8 | 1.86 |
| **#31** | 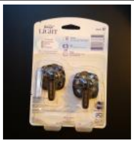 | 13.23 ± 0.49 | 2.67% ± 0.30% | 2 | Household goods packaging | 31.3 | 55.1 | 1.76 |
| **#32** | 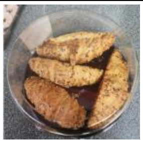 | 20.92 ± 0.2 | 2.71% ± 0.14% | 7 | Food packaging | 32.5 | 58.3 | 1.79 |

Supplementary Information Fig. 8 continued

| Sample number | Postconsumer Plastic products | Initial mass (mg) | Crystallinity % | Time for complete degradation (days) | Category | Mn kg/mol | Mw kg/mol | Đ |
|---|---|---|---|---|---|---|---|---|
| **#33** | 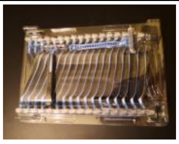 | 18.4 ± 0.1 | 2.90% ± 0.16% | 4 | Household goods packaging | 28.7 | 52.4 | 1.83 |
| **#34** | 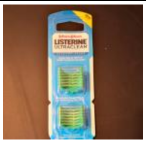 | 7.12 ± 1.16 | 2.93% ± 0.02% | 2 | Household goods packaging | 36.8 | 67.7 | 1.84 |
| **#35** | 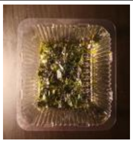 | 8.27 ± 1.84 | 3.00% ± 0.61% | 3 | Food packaging | 28.3 | 50.2 | 1.77 |
| **#36** | 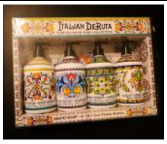 | 18.14 ± 0.13 | 3.06% ± 0.30% | 3 | Household goods packaging | 29.8 | 52.7 | 1.77 |
| **#37** | 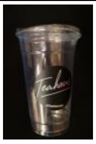 | 9.79 ± 0.06 | 3.21% ± 0.27% | 3 | Beverage packaging | 33.2 | 57.5 | 1.73 |
| **#38** | 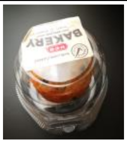 | 4.01 ± 0.54 | 3.42% ± 0.45% | 2 | Food packaging | 30.9 | 56.1 | 1.82 |
| **#39** | 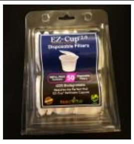 | 6.84 ± 0.45 | 3.47% ± 0.67% | 2.5 | Beverage packaging | 35.7 | 64.1 | 1.80 |
| **#40** | 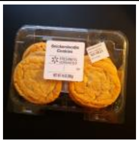 | 11.08 ± 0.77 | 3.56% ± 0.66% | 6 | Food packaging | 39.7 | 68.7 | 1.73 |
| **#41** | 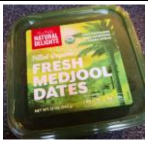 | 14.04 ± 0.4 | 3.57% ± 0.16% | 6 | Food packaging | 31.8 | 57.0 | 1.79 |
| **#42** | 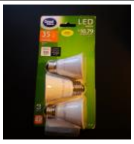 | 8.43 ± 0.48 | 3.58% ± 0.19% | 2.5 | Household goods packaging | 31.2 | 58.0 | 1.86 |
| **#43** | 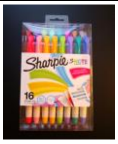 | 12.34 ± 0.3 | 3.72% ± 0.24% | 6 | Office supplies packaging | 36.0 | 66.5 | 1.85 |

Supplementary Information Fig. 8 continued

| Sample number | Postconsumer Plastic products | Initial mass (mg) | Crystallinity % | Time for complete degradation (days) | Category | Mn kg/mol | Mw kg/mol | Đ |
|---|---|---|---|---|---|---|---|---|
| **#44** | 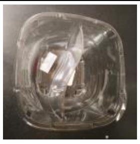 | 14.46 ± 0.43 | 4.09% ± 0.18% | 5 | Food packaging | 31.4 | 56.2 | 1.79 |
| **#45** | 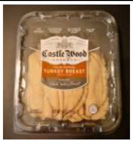 | 16.94 ± 0.12 | 4.56% ± 0.26% | 7 | Food packaging | 34.5 | 62.7 | 1.82 |
| **#46** | 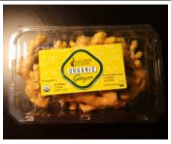 | 7.9 ± 2.78 | 4.69% ± 1.08% | 2 | Food packaging | 31.6 | 55.7 | 1.76 |
| **#47** | 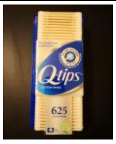 | 11.49 ± 0.2 | 4.85% ± 0.36% | 4 | Household goods packaging | 35.1 | 64.1 | 1.83 |
| **#48** | 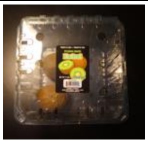 | 17.12 ± 0.16 | 4.85% ± 1.08% | 5.5 | Food packaging | 33.1 | 57.9 | 1.75 |
| **#49** | 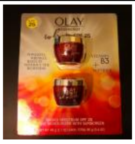 | 6.51 ± 2.65 | 5.30% ± 0.18% | 1.5 | Cosmetics packaging | 28.8 | 50.9 | 1.77 |
| **#50** | 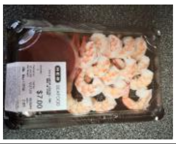 | 10.96 ± 0.21 | 5.79% ± 0.11% | 3.5 | Household goods packaging | 32.6 | 58.8 | 1.80 |
| **#51** | 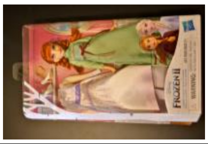 | 23.1 ± 0.05 | 6.24% ± 0.25% | 7 | Toy packaging | 32.2 | 59.8 | 1.86 |

Supplementary Information Fig. 8 continued

**a**

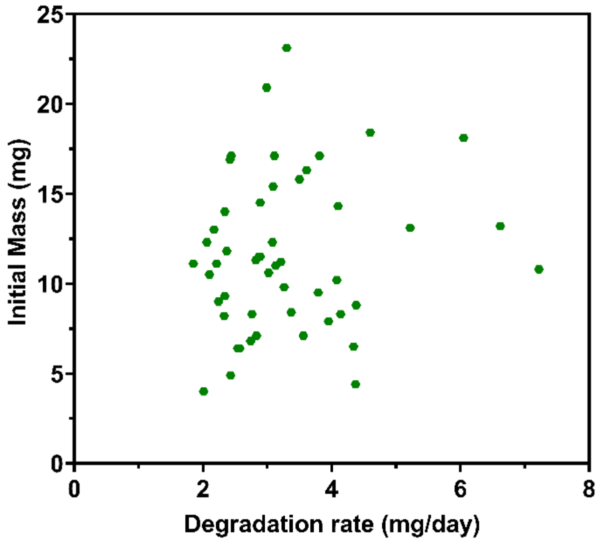
Initial Mass (mg)
Degradation rate (mg/day)
0
5
10
15
20
25
0
2
4
6
8

**b**

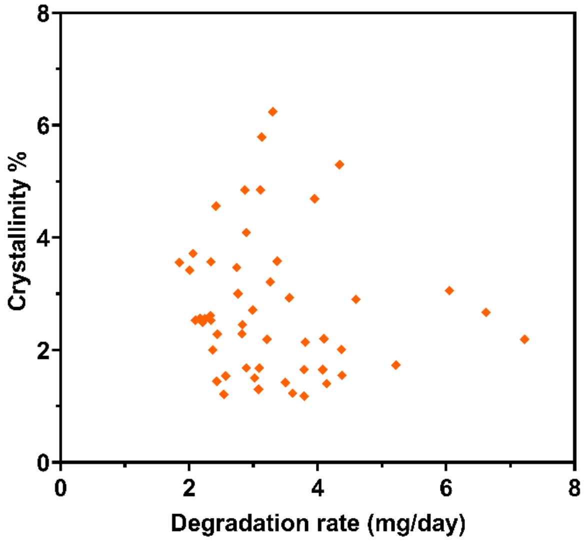
Crystallinity %
Degradation rate (mg/day)
0
2
4
6
8
0
2
4
6
8

**c**

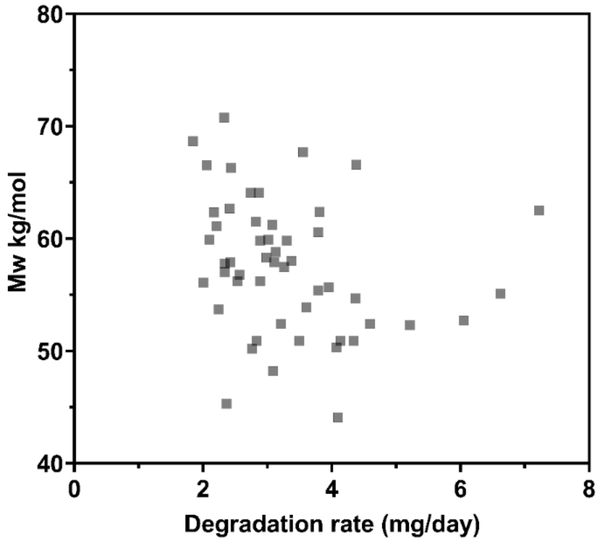
Mw kg/mol
Degradation rate (mg/day)
40
50
60
70
80
0
2
4
6
8

**d**

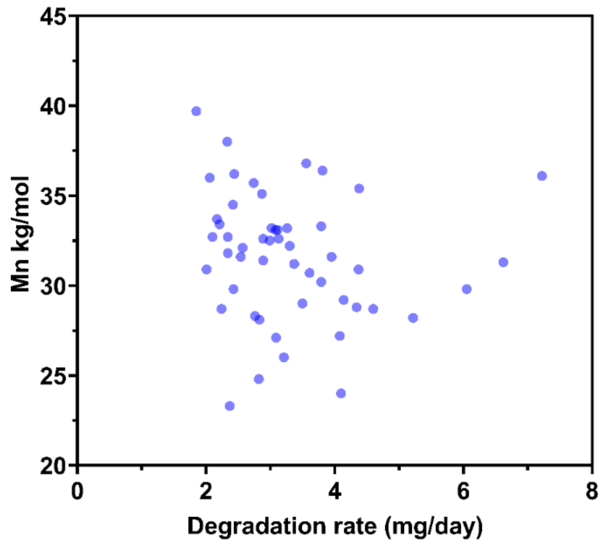


**e**

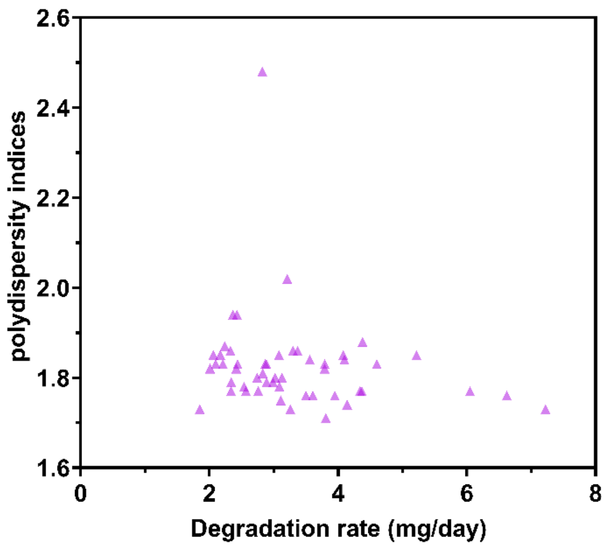


**Supplementary Information Fig. 9 | Scatterplot of degradation rate versus (a.) initial mass or (b.) crystallinity % or (c.) weight average molecular weight (Mw) or (d.) number average molecular weight (Mn) or (e.) polydispersity indices of the hole-punched films from 51 different post-consumer plastic products**. Degradation rate was not found to be dependent on any one metric of these various plastics.

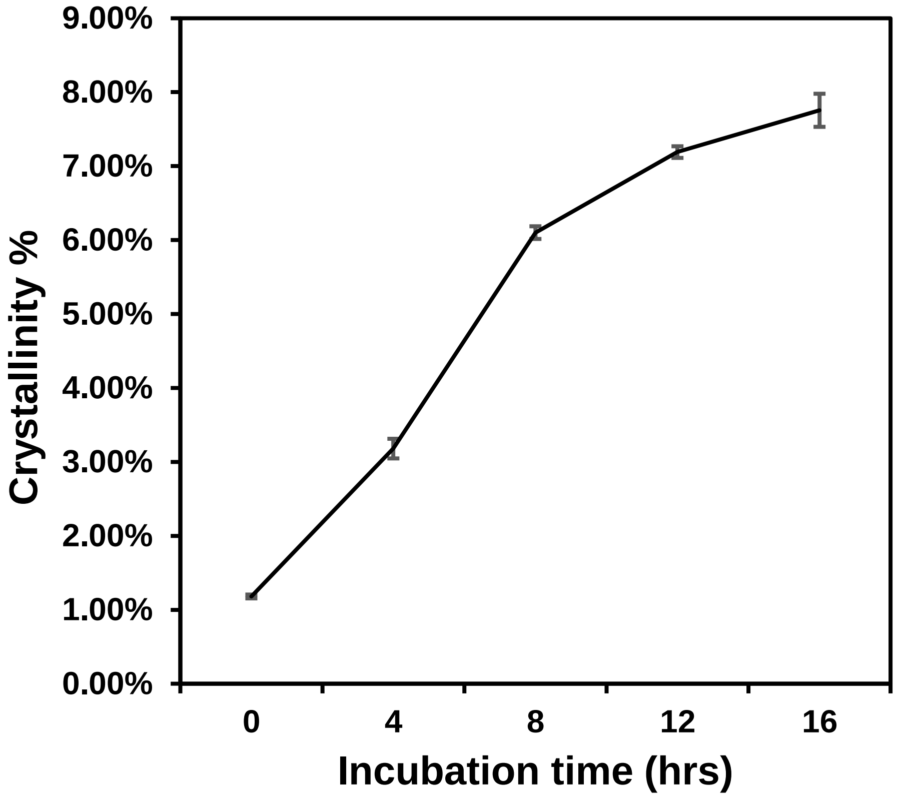


**Supplementary Information Fig. 10 | Time-course of crystallinity % of the degraded pc-PET film.** The hole-punched PET films from a bean cake PET container were treated with FAST-PETase for 0 hr, 4hr, 8 hr,12 hrs 16 hr in 100 mM $KH_2PO_4$-NaOH (pH 8.0) buffer at 50 ºC. Crystallinity % of the films was determined by DSC. All measurement were conducted in duplicate (n=2).

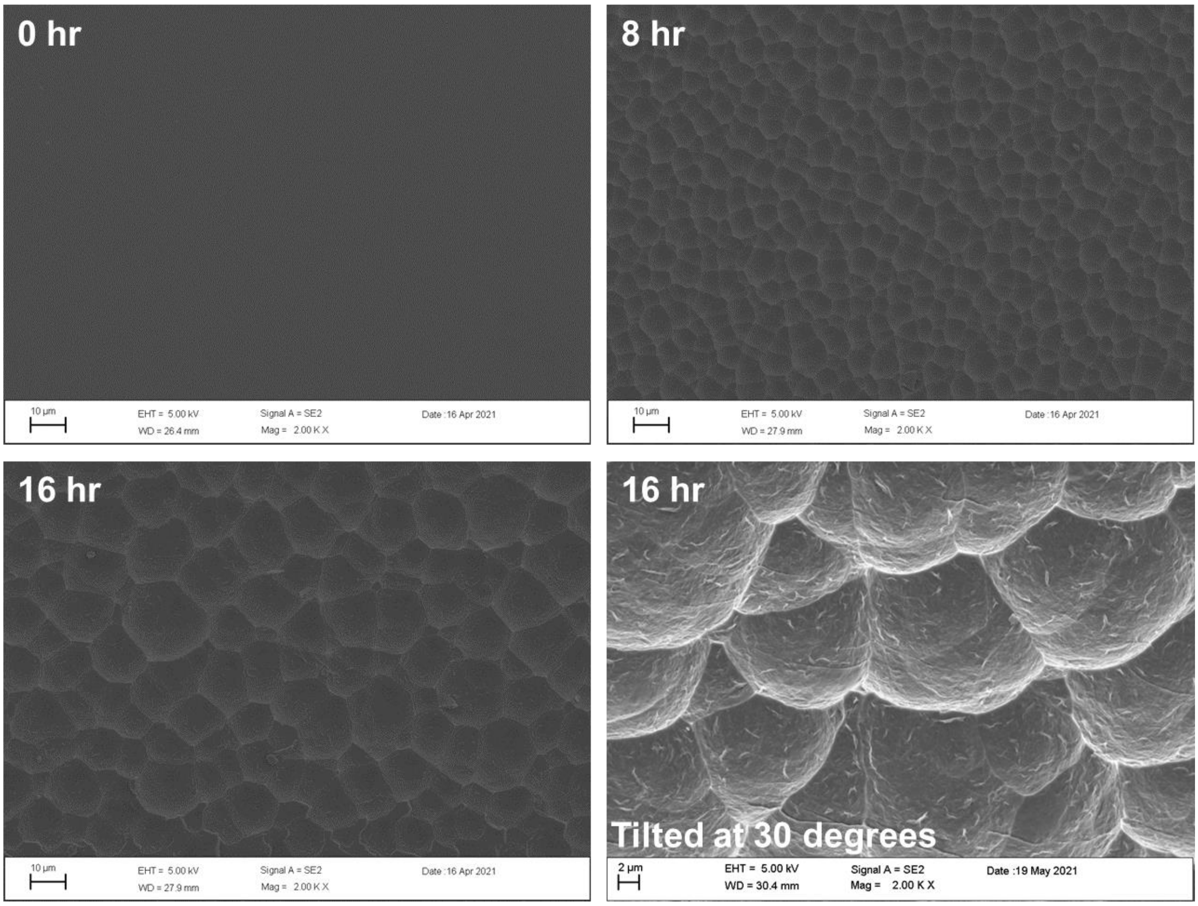


**Supplementary Information Fig. 11 | Scanning electron microscopic analysis of the pc-PET films.** The hole-punched PET films from a bean cake PET container were treated with FAST-PETase for 0 hr, 8 hr, 16 hr in 100 mM $KH_2PO_4$-NaOH (pH 8.0) buffer at 50 ºC.

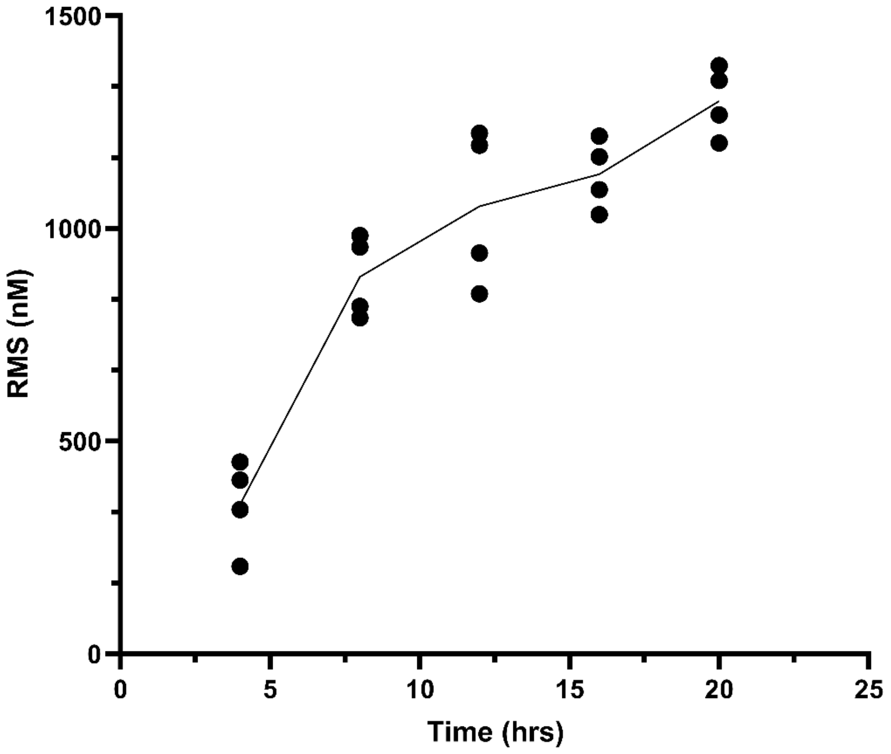


**Supplementary Information Fig. 12 | The surface roughness of the pc-PET films determined by atomic force microscopy.** The hole-punched PET films from a bean cake PET container were treated with FAST-PETase for 4 hr, 8 hr, 12 hr, 16 hr and 20 hr in 100mM $KH_2PO_4$-NaOH (pH 8.0) buffer at 50 ºC. The time-course profile of the surface roughness indicated that longer exposure times with FAST-PETase resulted in higher degree of surface roughness on the pc-PET films. RMS represents root mean square.

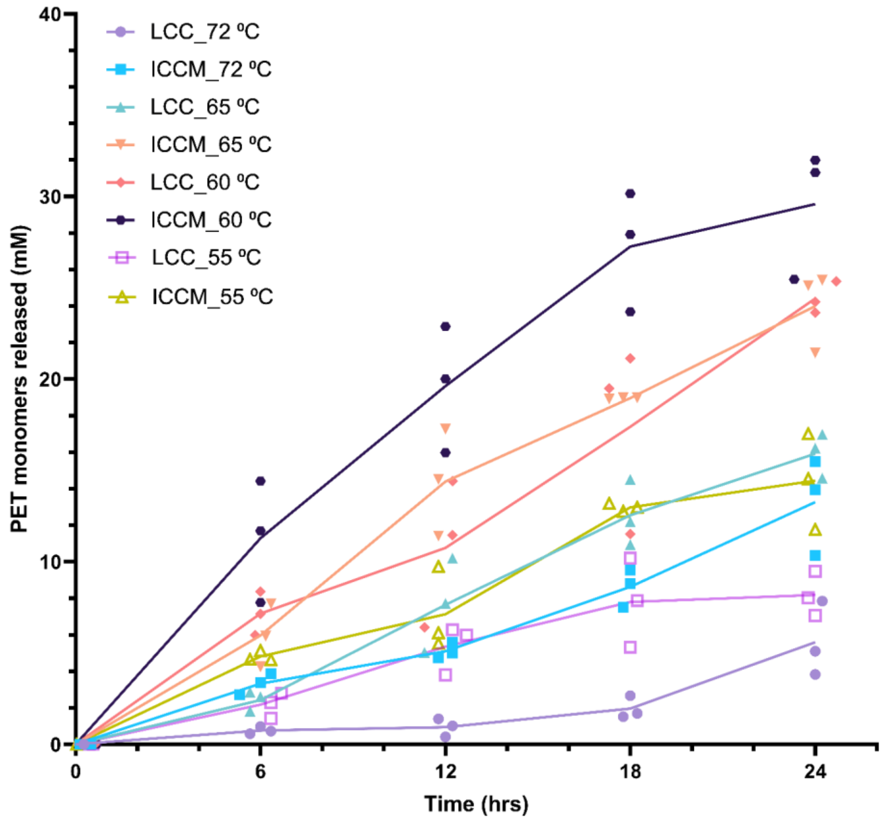


**Supplementary Information Fig. 13 | Time-course of PET-hydrolytic activity of LCC and ICCM at reaction temperatures of 55 ºC, 60 ºC, 65 ºC, and 72 ºC.** PET-hydrolytic activity was evaluated by measuring the amount of PET monomers (the sum of TPA and MHET) released from hydrolyzing the pc-PET (Bean cake plastic container) film by the tested PHEs at various time points. 100 mM $KH_2PO_4$-NaOH (pH 8.0) buffer was used for all reactions shown in this figure. All measurement were conducted in triplicate (n=3).

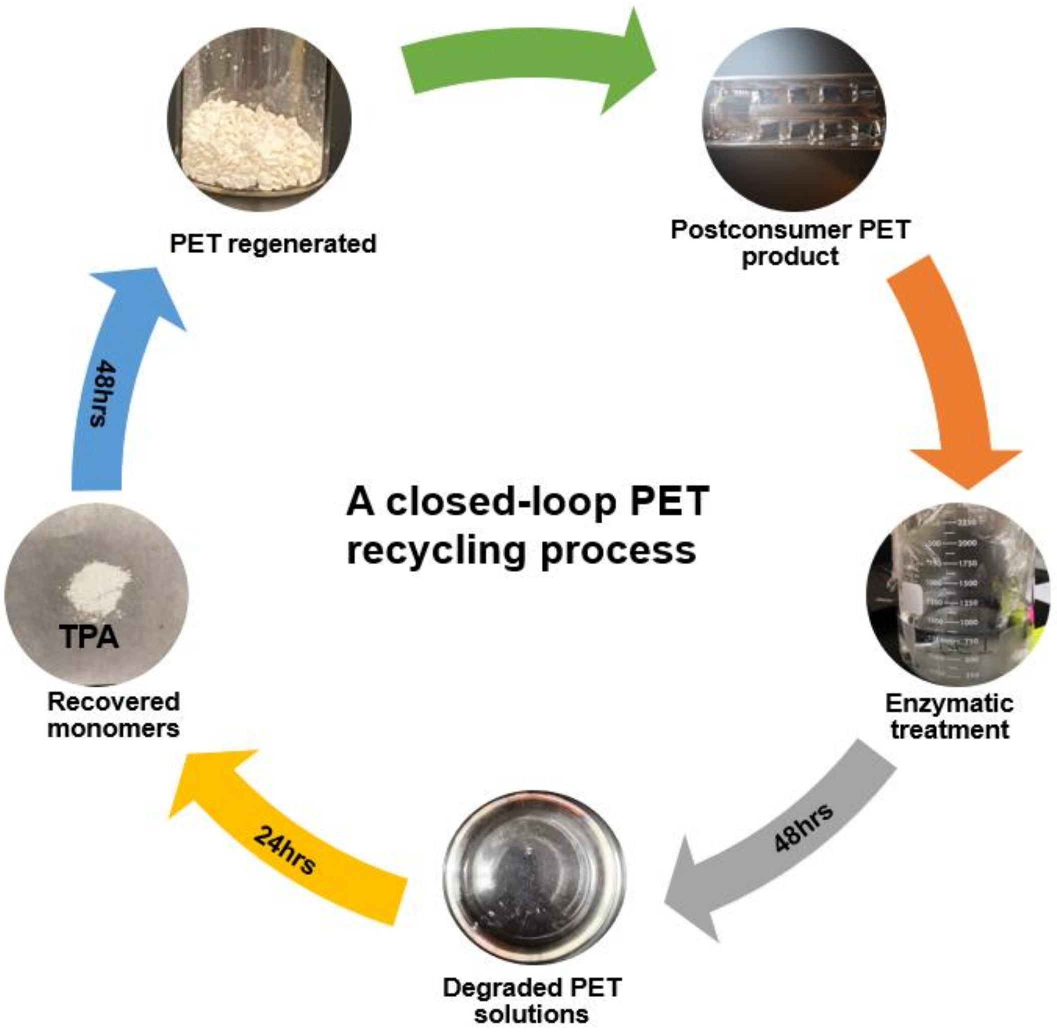


**Supplementary Information Fig. 14 | A closed-loop PET recycling process**. Demonstration of a closed-loop process for enzymatically degrading and then regenerating PET in the course of several days.

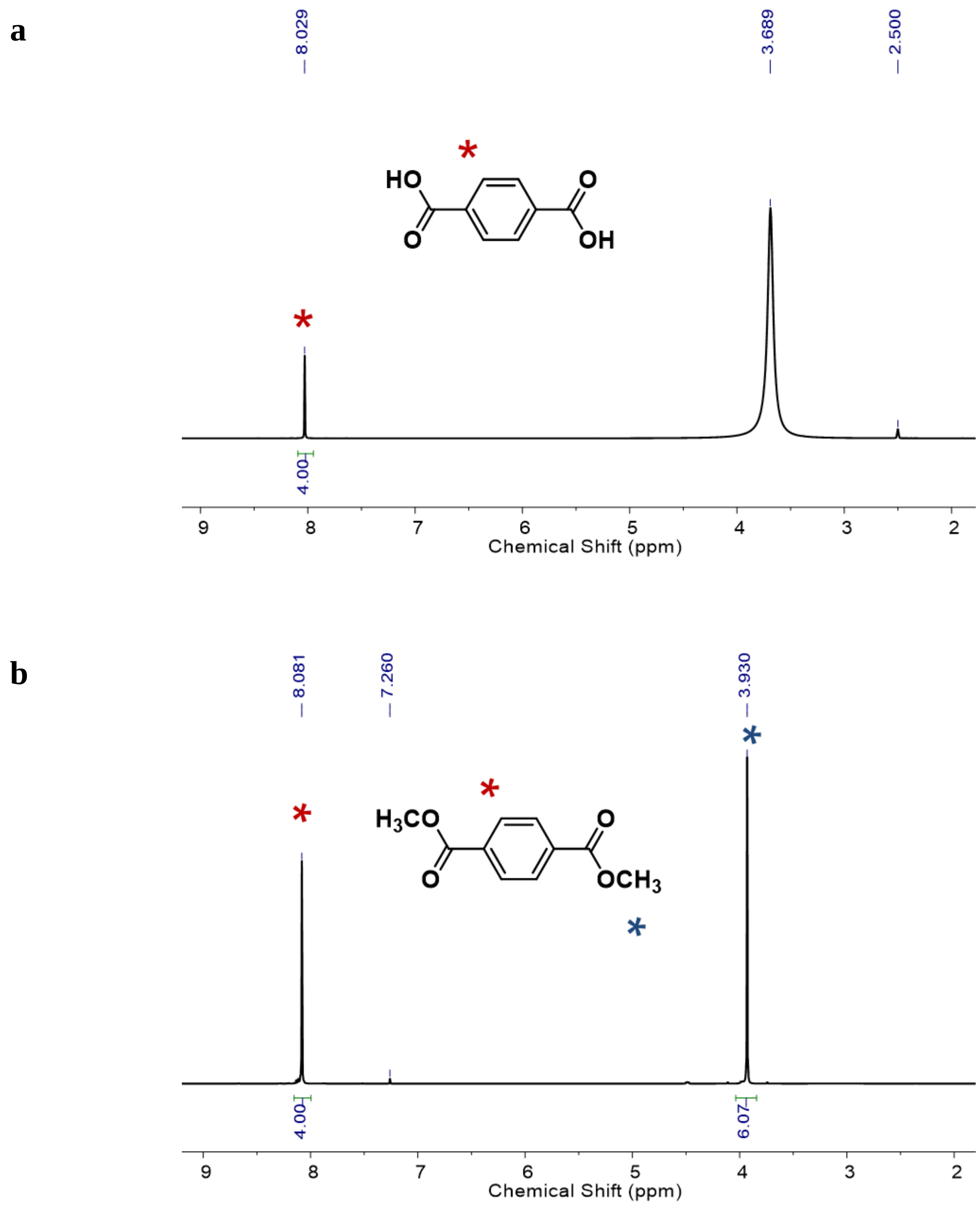


**Supplementary Information Fig. 15 | a. $^1$H NMR (400 MHz, $d_6$-DMSO) spectra of TPA recovered from degraded PET solutions.** The peak at 8.029 ppm corresponds to the hydrogen nuclei of the benzene ring. **b. $^1$H NMR (400 MHz, $CDCl_3$) spectra of DMT synthesized from TPA.** The peak at 8.081 ppm corresponds to the hydrogen nuclei of the benzene ring. The peak at 3.93 ppm corresponds to the hydrogen nuclei of the methyl group.

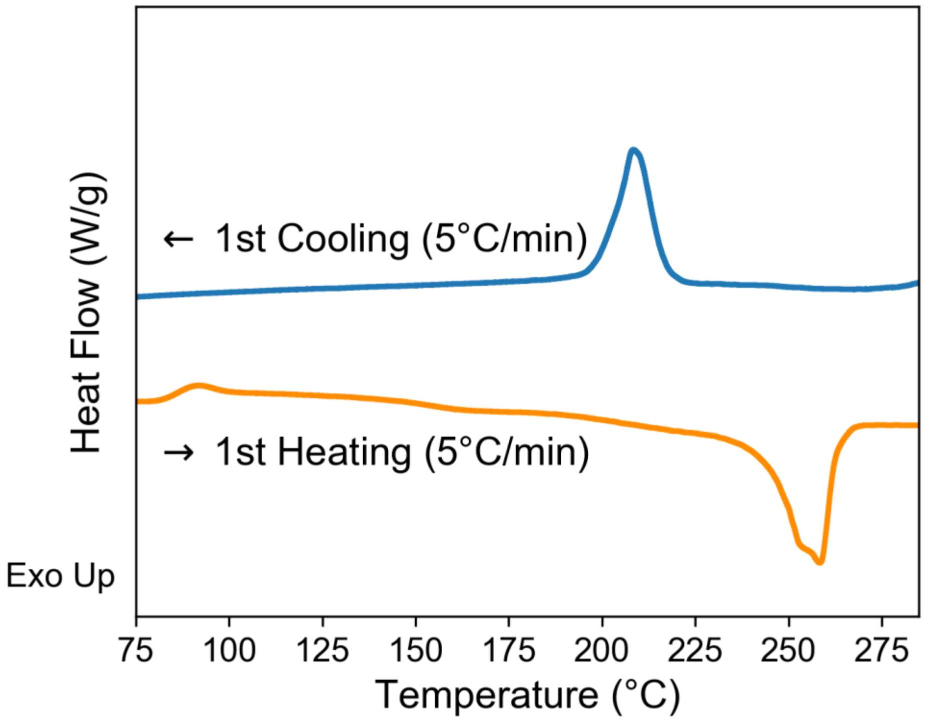


**Supplementary Information Fig. 16 | DSC trace of PET regenerated from the degraded solutions.** The crystallinity of this regenerated PET is 58.46%. The melting onset is 243.6 °C. The melting peak temperature is 258.4 °C. The glass transition temperature is 84.3 °C.

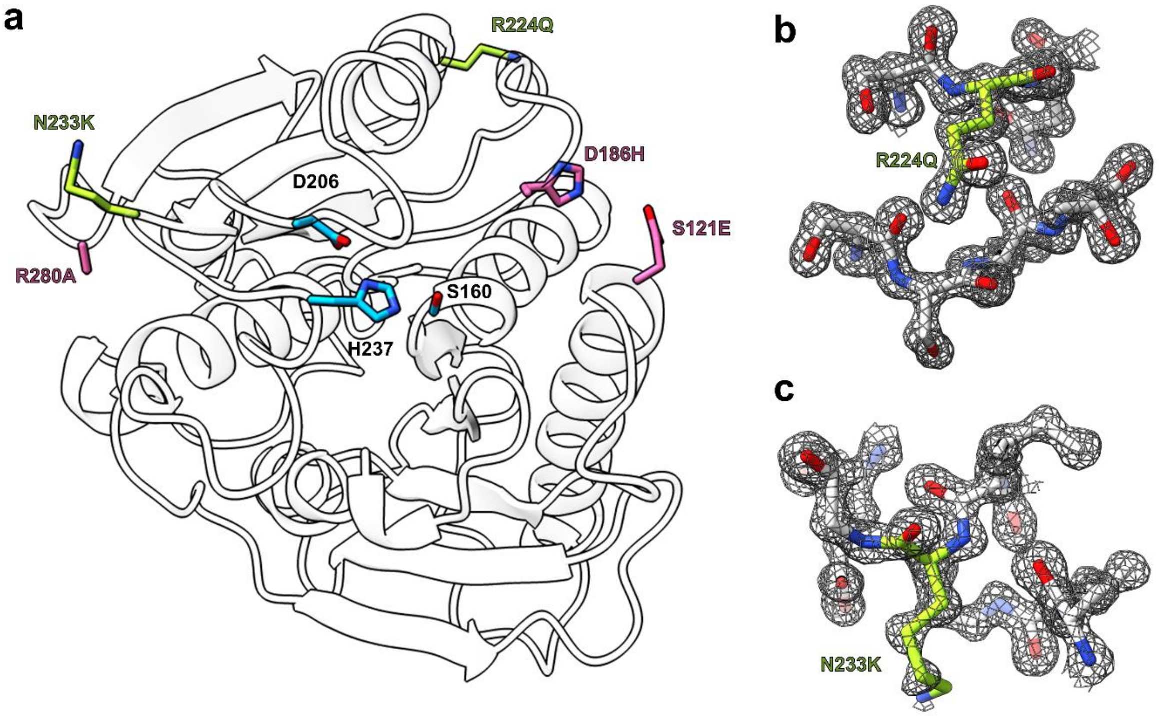


**Supplementary Information Fig. 17 | X-ray crystal structure of FAST-PETase. a.** Overall crystal structure of FAST-PETase. Catalytic triads (S160, D206, H237) are shown in blue sticks. Mutations originating from ThermoPETase (S121E, D186H, R280A) are shown in pink sticks, and novel mutations predicted by the neural network are shown in green-yellow sticks. **b-c.** $2F_o$-$F_c$ map (contoured at 1.5σ) shown as grey mesh superimposed on the stick models of novel mutation sites (**b.**) R224Q, (**c.**) N233K.

| **Data collection** | |
|---|---|
| Space group | $P2_12_12_1$ |
| Cell dimensions | |
| a, b, c (Å) | 50.9, 51.2, 84.1 |
| α, β, γ (°) | 90.0, 90.0, 90.0 |
| Resolution (Å) | 50.00-1.44 (1.46-1.44)* |
| $R_{sym}$/ $R_{pim}$ | 0.074(0.195)/0.031(0.113) |
| CC ½ $^{\Upsilon}$ | 0.988 (0.948) |
| I / σ | 27.4 (3.96) |
| Completeness (%) | 99.4 (94.3) |
| Redundancy | 6.6 (3.5) |
| **Refinement** | |
| Resolution (Å) | 43.714 - 1.439 (1.490 - 1.439) |
| No. reflections | 40270 (3753) |
| $R_{work}$ | 0.1515 (0.1641) |
| $R_{free}^{\pm}$ | 0.1657 (0.2118) |
| **No. atoms** | 2344 |
| Protein | 1981 |
| Ligand/ion | 5 |
| Water | 358 |
| **B-factors ($Å^2$)** | |
| Protein | 7.8 |
| Ligand/ion | 17.6 |
| Water | 23.2 |
| **R.m.s. deviations** | |
| Bond lengths (Å) | 0.012 |
| Bond angles (°) | 1.18 |
| **Ramachandran plot** | |
| Favored | 97.68% |
| Allowed | 2.32% |
| Outliers | 0.00% |
| **Molprobity score** | 1.16 / 97th percentile |

*Values for the corresponding parameters in the outermost shell in parenthesis.
$^{\Upsilon}CC_{1/2}$ is the Pearson correlation coefficient for a random half of the data; the two numbers represent the lowest and highest resolution shell, respectively.
$^{\pm}R_{free}$ is the $R_{work}$ calculated for about 10% of the reflections randomly selected and omitted from refinement.

**Supplementary Information Fig. 18 | Statistics of the crystal structural determination of FAST-PETase.**

Information about the obtained crystal structure is provided.

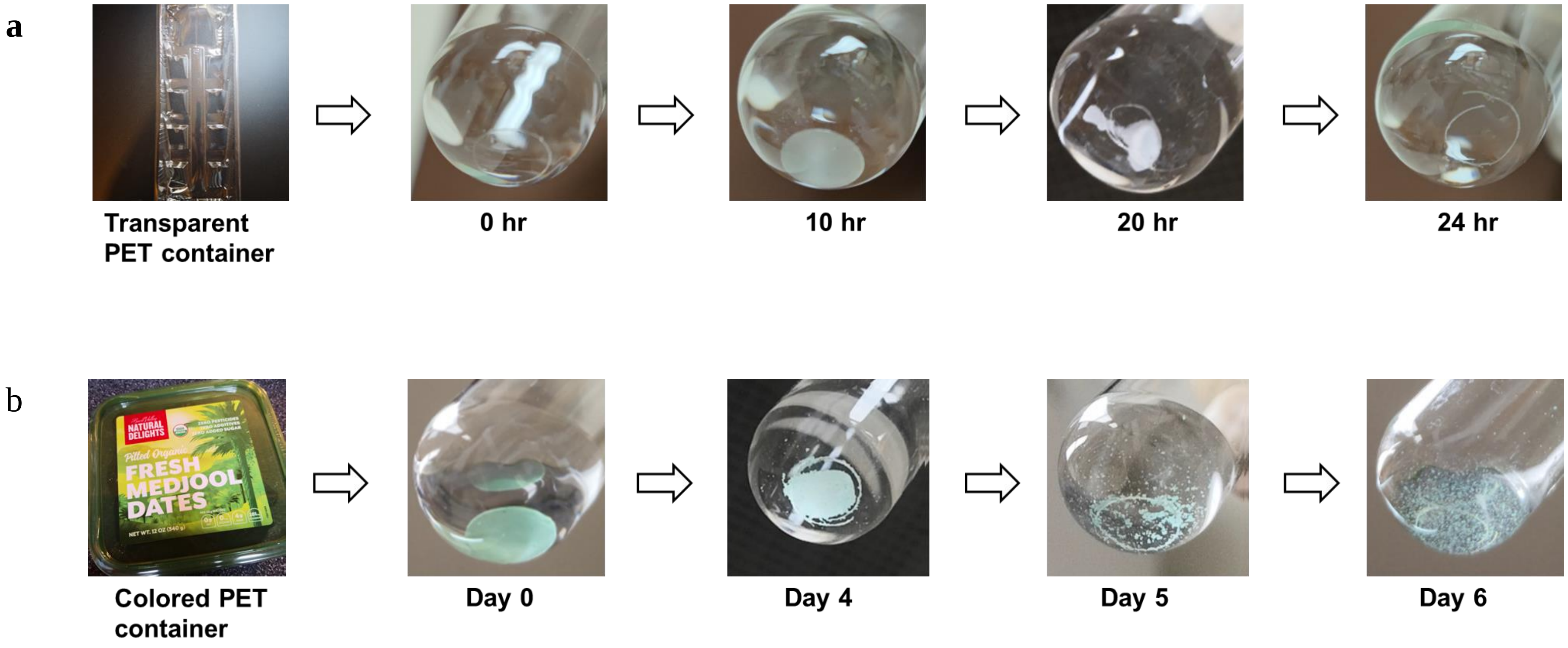


**Supplementary Information Fig. 19 | Stages of degradation of pc-PET films by FAST-PETase. a. The transparent pc-PET film (6 mm in diameter) was completely degraded (only cutting edges of the film remained) after 24 hrs of a single treatment with FAST-PETase at 50 $^{0}$C. b. The colored pc-PET film (6 mm in diameter) was completely degraded (only cutting edges of the film and some colorants remained) after six day of serial treatment with FAST-PETase at 50** $^{0}$C. Enzyme (200 nM) treatment was performed with 100 mM $KH_2PO_4$-NaOH (pH 8.0) buffer.
